# Targeting phosphodiesterase 10A disrupts MAPK signaling pathways in the tumor microenvironment to unleash antitumor immunity

**DOI:** 10.64898/2026.02.26.708383

**Authors:** Md Y. Gazi, Yan Ye, Ogacheko D. Okoko, Xin Wang, Samson E. Simon, Ahmad Alimadadi, Nandeeni Suryawanshi, Catlin Brandle, Mei Zheng, Lirong Pei, Khalda Fadlalla, Adam Keeton, Yulia Maxuitenko, Xi Chen, Austin W.T. Chiang, Patricia Schoenlein, Dongwen Lyu, David Wolff, Catherine C. Hedrick, Huidong Shi, Gary Piazza, Gang Zhou

## Abstract

Phosphodiesterase 10A (PDE10A), a cyclic nucleotide-degrading enzyme, is overexpressed in various human cancers. While PDE10A inhibition using small-molecule inhibitors or gene silencing suppresses tumor growth in xenograft models, its precise mechanism of action and immunological impact remain unclear. Here, we report that ADT-030, a novel PDE10A inhibitor, exhibits potent cytotoxicity against a broad range of murine tumor cell lines. ADT-030 is orally bioavailable and effectively suppresses tumor growth across multiple syngeneic mouse models. Notably, its efficacy is diminished in immunodeficient mice or upon CD8^+^ T cell depletion, highlighting a critical dependence on host immunity. The immunostimulatory properties of ADT-030 are further supported by its ability to induce immunogenic tumor cell death and promote dendritic cell (DC) maturation, its reliance on Batf3-expressing DCs to elicit antitumor CD8^+^ T cell response, and its synergy with anti-PD-1 therapy. Comprehensive immune profiling in the 4T1 breast cancer model, both in orthotopic and metastatic settings, revealed that ADT-030 selectively reduces myeloid-derived suppressor cells (MDSCs) while normalizing the immune landscape within the tumor. Mechanistically, ADT-030 disrupts multiple components of the mitogen-activated protein kinase (MAPK) signaling network in both tumor cells and MDSCs, leading to induction of apoptosis in these populations. These findings highlight the multi-faceted impact of PDE10A inhibition as a therapeutic strategy that not only disrupts tumor-intrinsic oncogenic signaling to inhibit tumor progression but also reshapes the tumor immune microenvironment to unleash antitumor immunity.

## Introduction

Phosphodiesterase 10A (PDE10A) is a dual-substrate phosphodiesterase that hydrolyzes both cyclic adenosine monophosphate (cAMP) and cyclic guanosine monophosphate (cGMP). PDE10A is overexpressed in multiple human cancers and plays a critical role in tumor cell proliferation and survival^1–3^. Notably, high PDE10A expression has been negatively correlated with overall and recurrence-free survival in non-small cell lung cancer (NSCLC) and ovarian cancer, supporting its potential as a therapeutic target^1,2,4–7^.

To develop PDE10A-targeted therapy suitable for oncology, we screened a chemically diverse library of sulindac derivatives to identify compounds that retain PDE10A inhibitory potency while eliminating cyclooxygenase (COX) inhibition to reduce toxicities associated with long-term sulindac administration. A first-generation derivative, ADT-061 (aka MCI-030), was identified as a selective PDE10A inhibitor that can effectively block oncogenic RAS signaling and Wnt/β-catenin transcriptional activity. ADT-061 demonstrated therapeutic efficacy in the *Apc*^+^/min-FCCC mouse model of colon tumorigenesis and in human ovarian cancer models^6,7^, establishing proof of concept for PDE10A inhibition as an anticancer strategy.

Both RAS and β-catenin signaling are critical drivers of cancer growth, metastasis, chemoresistance and immune evasion^8–10^. Gain of function mutations in RAS isoforms, including *KRAS*, *NRAS* and *HRAS*, occur across a wide range of human malignancies^11,12^. Aberrant RAS signaling in cancer cells, mediated primarily through the Raf-MEK-ERK (MAPK) and PI3K-AKT-mTOR cascades, is strongly associated with poor clinical outcomes^13^. In recent years, substantial progress has been made in targeting oncogenic RAS, culminating in FDA approval of the mutant-specific KRAS inhibitors sotorasib and adagrasib for the treatment of selected cancer types. These agents exhibit direct tumoricidal activity and, in preclinical models, have been shown to remodel the tumor immune microenvironment (TiME)^14^. However, their clinical utility is constrained by the rapid emergence of resistance and dose-limiting toxicities, which limit their use in combination regimens, including immune checkpoint blockade (ICB)^14,15^. To address these challenges, there is growing interest in developing broader-spectrum pan-KRAS and pan-RAS inhibitors capable of achieving more durable suppression of aberrant RAS signaling and overcoming resistance mechanisms^16–18^. Several such agents have advanced into clinical trials, underscoring the promise of this therapeutic strategy. Given that PDE10A inhibition suppresses multiple nodes within the RAS-MAPK signaling network, we hypothesized that PDE10 inhibitors may function as indirect pan-RAS pathway modulators, providing an alternative approach for treating malignancies driven by aberrant RAS signaling. Although we have previously shown that PDE10A inhibition suppresses RAS and β-catenin signaling through accumulation of cGMP and subsequent activation of protein kinase G (PKG)^1,2,5–7^, these studies were largely conducted using human cancer cell lines and xenograft models, leaving the immunological consequences of PDE10A inhibition largely unexplored.

Here, we report preclinical studies on ADT-030, a second-generation PDE10A inhibitor with improved oral bioavailability and optimized drug-like properties, across a range of syngeneic murine tumor models. Using the 4T1 triple-negative breast cancer (TNBC) model, we provide a comprehensive analysis on the immunological impact of ADT-030 in both orthotopic and metastatic settings. Our study uncovers the cellular and molecular mechanisms underlying ADT-030’s antitumor activity, demonstrating its ability to disrupt tumor-intrinsic oncogenic signaling, enhance tumor immunogenicity, normalize the TiME, and synergize with ICB. Collectively, these findings underscore the translational potential of PDE10A inhibition as both a standalone therapeutic strategy and a component of rational combination regimens integrated with targeted therapies or immunotherapies.

## Results

### ADT-030 is a highly selective PDE10A inhibitor that exhibits cytotoxic activity across murine tumor cell lines

ADT-030 is a novel, potent, and selective inhibitor of phosphodiesterase 10A (PDE10A), an enzyme that catalyzes the hydrolysis of cyclic guanosine monophosphate (cGMP) and cyclic adenosine monophosphate (cAMP) (manuscript in preparation). Our recent study demonstrates that ADT-030 exhibits single-agent antitumor activity in human pancreatic cancer cell lines and patient-derived xenograft models by suppressing oncogenic RAS/MAPK and β-catenin signaling pathways^19^. In the present study, we sought to evaluate the immunological impact of ADT-030 using syngeneic mouse tumor models. ADT-030 inhibited PDE10A enzymatic activity with an IC_50_ of 1.1 µM, determined by a cell-free assay measuring cGMP hydrolysis by recombinant PDE10A (Fig. 1A). As the intended molecular target, PDE10A protein was readily detected across a panel of murine tumor cell lines, including 4T1, CT26, MC38, A20, KPC, EL4, and B16 (Fig. 1B). ADT-030 exhibited broad cytotoxic activity across all tumor cell lines tested, with IC₅₀ values for growth inhibition closely aligning with the IC₅₀ observed for recombinant PDE10A inhibition (Fig. 1C). These findings suggest that PDE10 inhibition by ADT-030 may contribute to its cytotoxic effects on tumor cells.

**Fig. 1.**
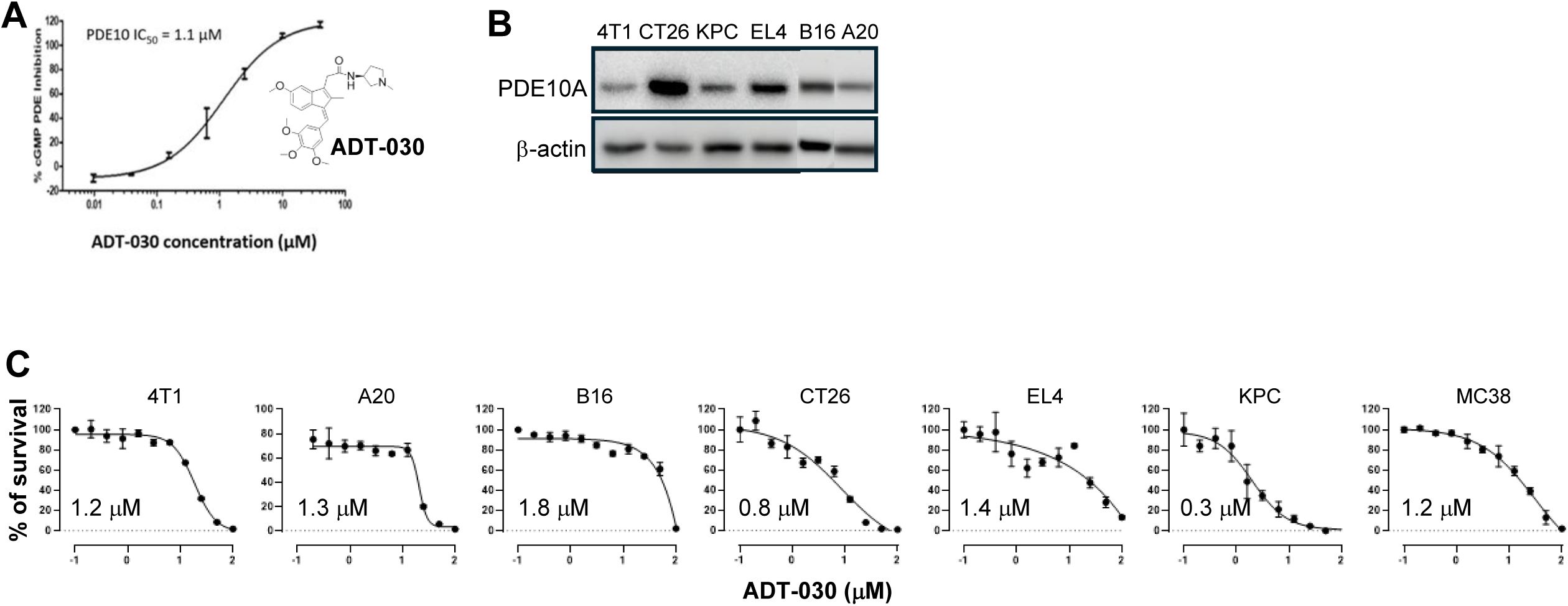
ADT-030 is a novel PDE10 inhibitor with broad cytotoxic activity against mouse tumor cell lines. (A) ADT-030 inhibits cGMP hydrolysis by recombinant PDE10 with an IC₅₀ of 1.1 μM. The chemical structure of ADT-030 is provided. (B) Western blot analysis showing PDE10A expression levels in selected mouse tumor cell lines, with β-actin used as loading control. (C) ADT-030–mediated cytotoxicity in tumor cells assessed by cell viability assays, with calculated IC₅₀ values indicated.

### ADT-030 treatment disrupts RAS signaling and induces apoptosis in tumor cells

We previously reported that ADT-007, an analog of ADT-030, inhibits oncogenic MAPK/Akt signaling in human colorectal and pancreatic cancer cell lines and suppresses the growth of RAS-active tumors *in vivo*^20^. Our recent work provides direct evidence that ADT-030 inhibits RAS/MAPK signaling in human pancreatic cancer cell lines and patient-derived xenografts^19^. These findings prompted us to examine the impact of ADT-030 on RAS signaling in syngeneic murine tumor models. As shown in Fig. 2A, basal RAS-GTP levels, assessed by a RAS-RBD pulldown assay, were detectable across all tumor cell lines examined and were, as expected, higher in CT26, EL4, and KPC cells, which harbor oncogenic *KRAS* mutations^21^. We next evaluated the effects of ADT-030 on downstream RAS signaling. Western blot (WB) analysis revealed reduced phosphorylation of cRaf (p-cRaf) and Akt (p-Akt) in ADT-030-treated 4T1 cells (Fig. 2B), indicating suppression of the RAS–PI3K–Akt–mTOR pathway. In parallel, ADT-030 treatment decreased phosphorylated ERK (p-ERK) while inducing activation of stress-associated MAPK pathways, including phosphorylated JNK/c-JUN (p-JNK/p–cJUN) and cleaved caspase-3 (cCas3) (Fig. 2C), consistent with engagement of pro-apoptotic signaling. Some of these molecular changes were further validated by flow cytometry, which revealed significant reductions in p-Akt and p-S6 accompanied by increased cCas3 in ADT-030-treated 4T1 cells (Fig. 2D). Similar signaling alterations, including reduced p-Akt, p-S6, and p-ERK, and increased p-JNK/p-cJUN, p-p53, and cCas3, were observed across multiple tumor cell lines following ADT-030 treatment (Supplemental Fig. 1). To define the transcriptional consequences of ADT-030–mediated signaling disruption, we performed bulk RNA sequencing on 4T1 cells treated with ADT-030 or DMSO for 16 hours. ADT-030 treatment caused profound transcriptomic changes in 4T1 cells, with large numbers of both upregulated and downregulated genes (Fig. 2E). Notably, many genes downregulated are associated with tumor proliferation, metastasis, angiogenesis, chemoresistance, and immune suppression, including *Igf1, Pdgfrb, Zeb1, Rab27a, Dusp6, Myc, Spp1, Dll4, Notch3, Wnt10a, Fzd4, Arg1, Il6,* and *Nt5e* (Fig. 2F). Among these, *Wnt10a* and *Fzd4* are components of the Wnt/β-catenin pathway, whereas *Dusp6* and *Myc* are known transcriptional targets of RAS signaling, consistent with our previous reports showing that PDE10 inhibition suppresses both RAS and β-catenin signaling. In contrast, ADT-030 treatment upregulated the genes linked to tumor suppression, cellular stress responses, and pro-inflammatory signaling, such as *Irf5, Irf6, Sall1, Atf3, Cxcl1,* and *Cxcl10* (Fig. 2F). Pathway enrichment analysis (Supplemental Fig. 2) revealed downregulation of receptor tyrosine kinase signaling, collagen formation, and ECM-receptor interaction pathways, which are pathways involved in supporting tumor growth, invasion, and stromal remodeling. In contrast, upregulated pathways included calcium signaling, DNA damage response, and apoptotic pathways, which are linked to cellular stress and programmed cell death. Collectively, these data support the hypothesis that ADT-030 suppresses tumor-intrinsic oncogenic programs, including RAS and Wnt/β-catenin pathways, while activating stress-associated MAPK pathways and immune-related transcriptional programs, which promote tumor cell apoptosis and enhance antitumor immunity.

**Fig. 2.**
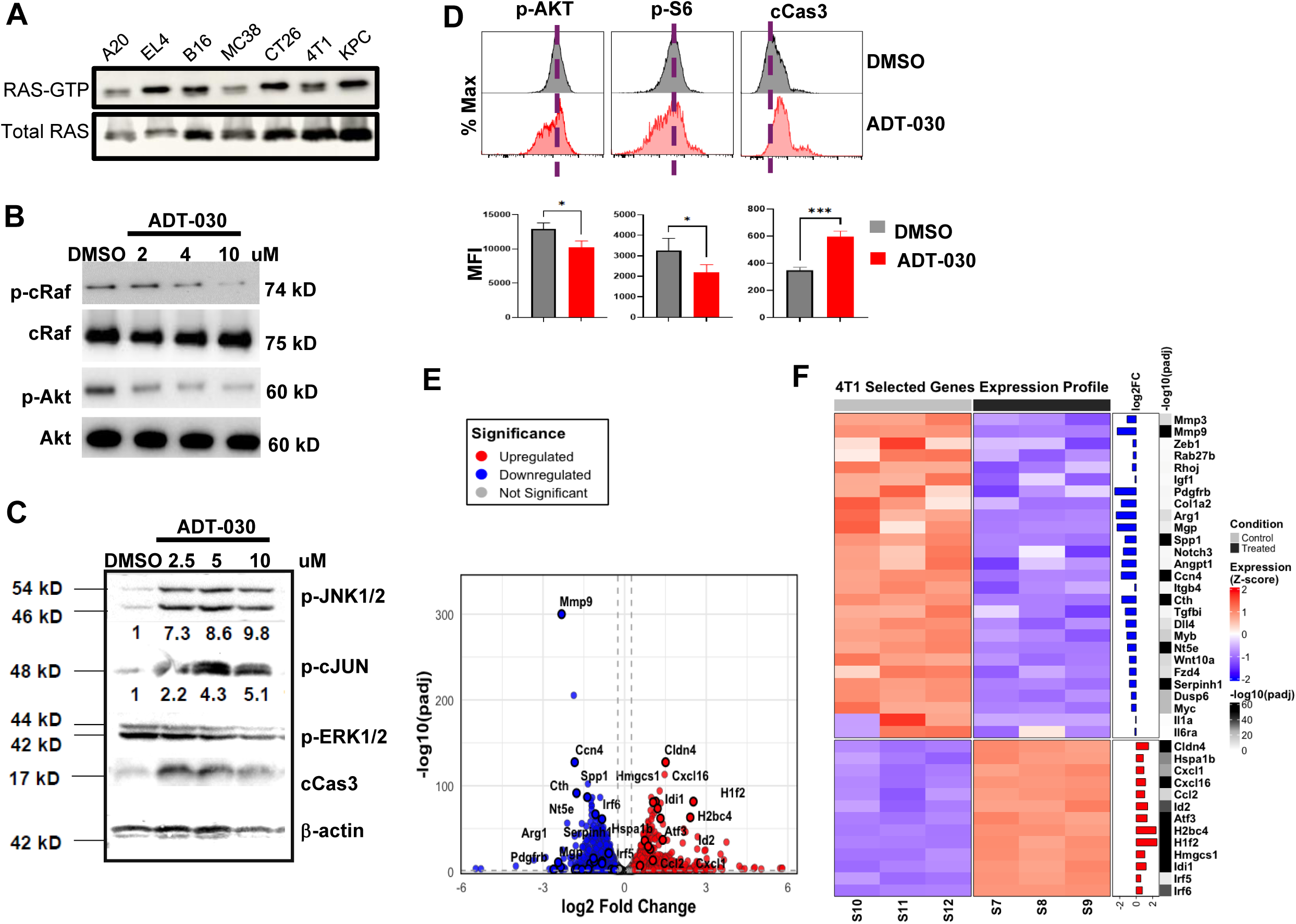
ADT-030 suppresses MAPK signaling in tumor cells. (A) RAS activity measured by RAS pull-down assay. Western blot shows GTP-bound RAS in tumor cells, with signal normalized to total RAS. (B) WB analysis of phosphorylated cRaf and AKT in 4T1 cells following ADT-030 treatment. Total cRaf and AKT are shown as loading controls. (C) WB analysis of phosphorylated JNK, c-JUN, ERK1/2, and cleaved caspase-3 in 4T1 cells after ADT-030 treatment. β-actin serves as a loading control. (D) Representative flow cytometry profiles of p-AKT, p-S6, and cleaved caspase-3 in DMSO- and ADT-030–treated 4T1 cells. Mean fluorescence intensities (MFI) are summarized in bar graphs. *, *P* < 0.05; ***, *P* < 0.001. (E) Volcano plot of bulk RNA-seq analysis of 4T1 cells treated with ADT-030 or DMSO for 16h. Each dot represents one gene. Red dots indicate upregulated genes, and blue dots indicate downregulated genes. (F) Heat map of selected differentially expressed genes in ADT-030-treated versus control 4T1 cells. Genes shown include regulators of tumor growth, RAS and Wnt signaling, and immune-related pathways..

### ADT-030 is orally bioavailable and mediates tumor growth inhibition in vivo across multiple murine tumor models

We next evaluated whether the potent cytotoxic effects of ADT-030 observed *in vitro* could be recapitulated *in vivo*. Tumor-bearing mice were treated once daily with vehicle or ADT-030 by oral gavage beginning when tumors became palpable (Fig. 3A schema). A pilot dose-escalation study in the 4T1 model demonstrated that daily oral administration of ADT-030 at 100-150 mg/kg was well tolerated and conferred significant therapeutic benefit (data not shown). Based on these findings, 150 mg/kg was selected for subsequent experiments.

**Fig. 3.**
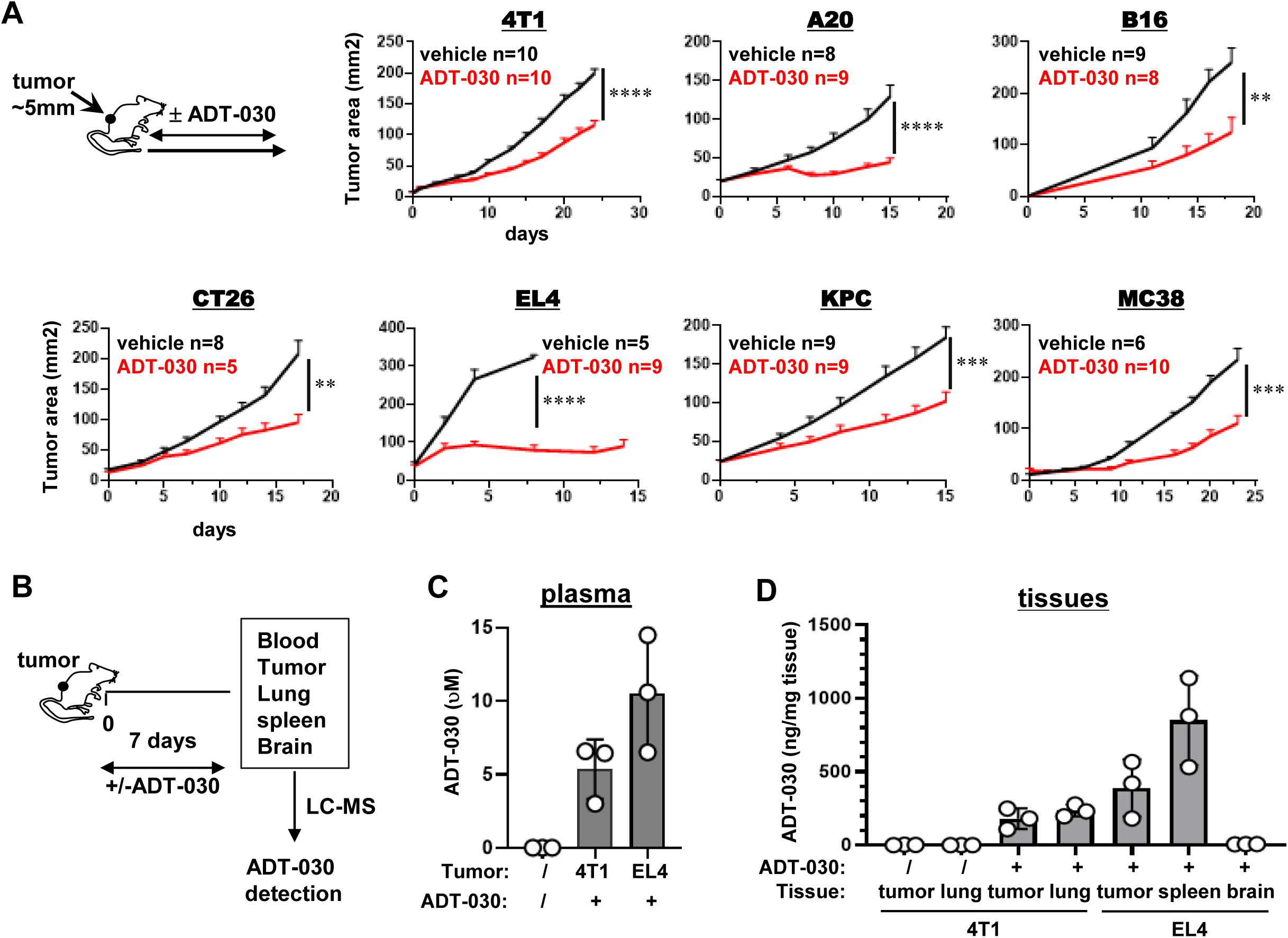
ADT-030 demonstrates in vivo efficacy across multiple mouse tumor models. The experimental timeline is shown in the schema. 4T1 cells were orthotopically implanted into the mammary fat pads of female mice, while other tumor cell lines were implanted subcutaneously into the rear flanks. When tumors became palpable (∼5 mm), mice were treated daily by oral gavage with vehicle or ADT-030 (150 mg/kg). (A) Tumor growth curves for each treatment group across tumor models. Data are shown as mean ± SEM of tumor area over time after treatment with number of mice in each group indicated. **, *P* < 0.01; ***, *P* < 0.001; ****, *P* < 0.0001. (B) Schematic of the timeline for plasma and tissue collection for LC–MS quantification of ADT-030. (C) Plasma concentrations of ADT-030 in tumor-bearing mice. Plasma from untreated naïve mice serves as a control. Data are shown as mean ± SEM. (D) ADT-030 concentrations in the indicated organs. Tissues from untreated 4T1-bearing mice were included as controls. Data are shown as mean ± SEM.

As shown in Fig. 3A, ADT-030 treatment significantly delayed tumor growth across all seven murine tumor models tested, with EL4 tumors exhibiting the greatest sensitivity. Importantly, mice maintained stable body weight throughout treatment, as demonstrated in multiple tumor models (Supplemental Fig. 3A). To further assess potential toxicity, histological analyses were performed on major organs harvested from tumor-free naïve mice following a 10-day course of vehicle or ADT-030 treatment. H&E staining revealed no gross pathological abnormalities in the heart, brain, lung, liver, kidney, or spleen (Supplemental Fig. 3B), indicating that ADT-030 does not induce detectable organ toxicity at the tested dose.

To evaluate drug bioavailability and tissue distribution, we selected the EL4 and 4T1 tumor models for pharmacokinetic analysis. Plasma and organs were collected from vehicle- or ADT-030-treated tumor-bearing mice and analyzed by LC/MS (Fig. 3B). Plasma concentrations of ADT-030 ranged from 5 to 10 μM (Fig. 3C), exceeding the reported IC₅₀ (∼1 μM) for PDE10A inhibition. In EL4-bearing mice, ADT-030 was readily detectable in tumors and spleens but was largely absent from the brain (Fig. 3D). It is important to note that limited central nervous system exposure is a desirable safety feature of ADT-030 given the high PDE10A expression in the brain^22^. 4T1 is a well-established model of triple-negative breast cancer (TNBC) in which orthotopically implanted tumor cells preferentially metastasize to the lungs^23^. In 4T1-bearing mice, ADT-030 accumulated in both primary tumors and lungs (Fig. 3D), consistent with activity at both primary and metastatic sites. Collectively, these findings establish ADT-030 as an orally bioavailable, efficacious, and well-tolerated therapeutic candidate with broad antitumor activity *in vivo*.

### ADT-030 reduces metastatic burden in the 4T1 lung metastasis model

Metastatic disease remains the leading cause of cancer-related mortality, including in TNBC. Given the ability of ADT-030 to accumulate in the lungs, we investigated its effects on metastatic tumor growth using a 4T1 experimental lung metastasis model. As depicted in Fig. 4A schema, luciferase-expressing 4T1 cells (4T1.luci) were intravenously (i.v.) injected into mice, followed by initiation of vehicle or ADT-030 treatment 3 days later and continued for 28 days. Bioluminescence imaging (BLI) revealed a significant reduction in lung tumor burden in ADT-030-treated mice compared with vehicle-treated controls (Fig. 4A). Consistent with this, ADT-030 treatment significantly prolonged mouse survival in this aggressive metastasis model (Fig. 4B). Notably, the therapeutic benefit of ADT-030 was evident when treatment was initiated within 3 days of tumor injection, whereas delaying treatment until day 10 diminished efficacy (Supplemental Fig. 4). These results suggest that ADT-030 monotherapy is more effective at disrupting early metastatic colonization than at treating established metastatic lesions. To directly assess the impact of ADT-030 on tumor colonization, mice injected i.v. with 4T1.luci cells were treated with vehicle or ADT-030 starting on the day of tumor inoculation (Fig. 4C). Immediate in vivo and ex vivo BLI confirmed comparable tumor cell seeding in the lungs of both groups shortly after injection (Fig. 4D). However, seven days later, vehicle-treated mice exhibited aggressive lung tumor growth, whereas ADT-030–treated mice showed a marked reduction in tumor burden (Fig. 4E). Together, these findings demonstrate that ADT-030 interferes with early metastatic colonization, highlighting its potential as a therapeutic agent for the treatment of metastatic disease.

**Fig. 4.**
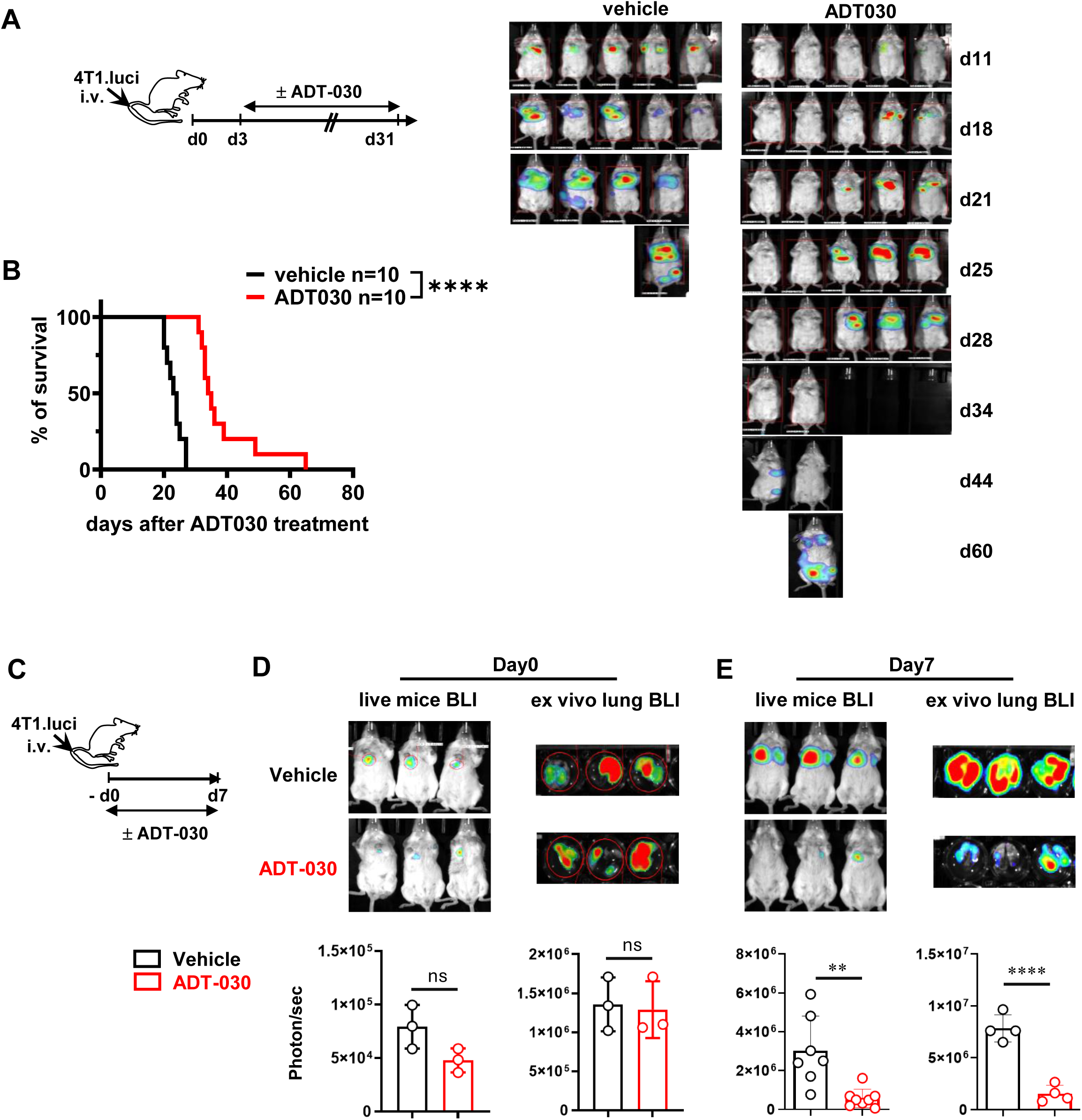
ADT-030 inhibits the colonization and growth of metastatic 4T1 tumors. (A) ADT-030 prolongs survival in an experimental 4T1 metastasis model. Schematic shows the treatment timeline. In brief, mice injected with 4T1.luci cells via i.v. were treated with either vehicle or ADT-030 for 21 days, starting 3 days after tumor inoculation. Representative BLI showing tumor burden at the indicated time points is shown. Kaplan-Meier survival curves are shown in (B). (C) Schema depicting the experimental design to assess the effect of ADT-030 on tumor colonization. Mice were injected intravenously with 4T1.luci cells and treated with vehicle or ADT-030 beginning shortly after tumor inoculation. Tumor burden in live mice and in isolated lungs was quantified by BLI. (D, E) Representative BLI images of tumor signal in mice and lungs acquired 2 h after tumor inoculation (D) and after 7 days of treatment (E). Quantification of photon flux (photons/s) is shown as mean ± SEM in bar graphs.. ns, not significant; **, *P* < 0.01; ****, *P* < 0.0001.

### ADT-030-mediated tumor growth inhibition is immune dependent and enhances anti-PD-1 therapy

To determine whether the antitumor activity of ADT-030 requires host immunity, we evaluated its efficacy in immunodeficient mouse models. Across multiple tumor models, the antitumor effects of ADT-030 were markedly diminished in immunodeficient NSG mice (Fig. 5A) and in RAG2-deficient mice (Supplemental Fig. 5), indicating a requirement for adaptive immunity. Consistent with this observation, depletion of CD8⁺ T cells significantly impaired the antitumor efficacy of ADT-030 in immunocompetent mice in multiple tumor models (Fig. 5B), underscoring a critical role for endogenous CD8⁺ T cells in mediating ADT-030-induced antitumor immunity. Given its immune-dependent activity, we next evaluated whether ADT-030 could enhance the efficacy of immune checkpoint blockade. In mice bearing subcutaneous CT26 or MC38 tumors, combination treatment with ADT-030 and anti-PD-1 antibody markedly improved survival compared with either monotherapy alone (Fig. 5C). These findings demonstrate that ADT-030 not only has the capacity to elicit host antitumor immunity but also can potentiate the therapeutic efficacy of PD-1 blockade, supporting its potential utility in combination immunotherapy strategies.

**Fig. 5.**
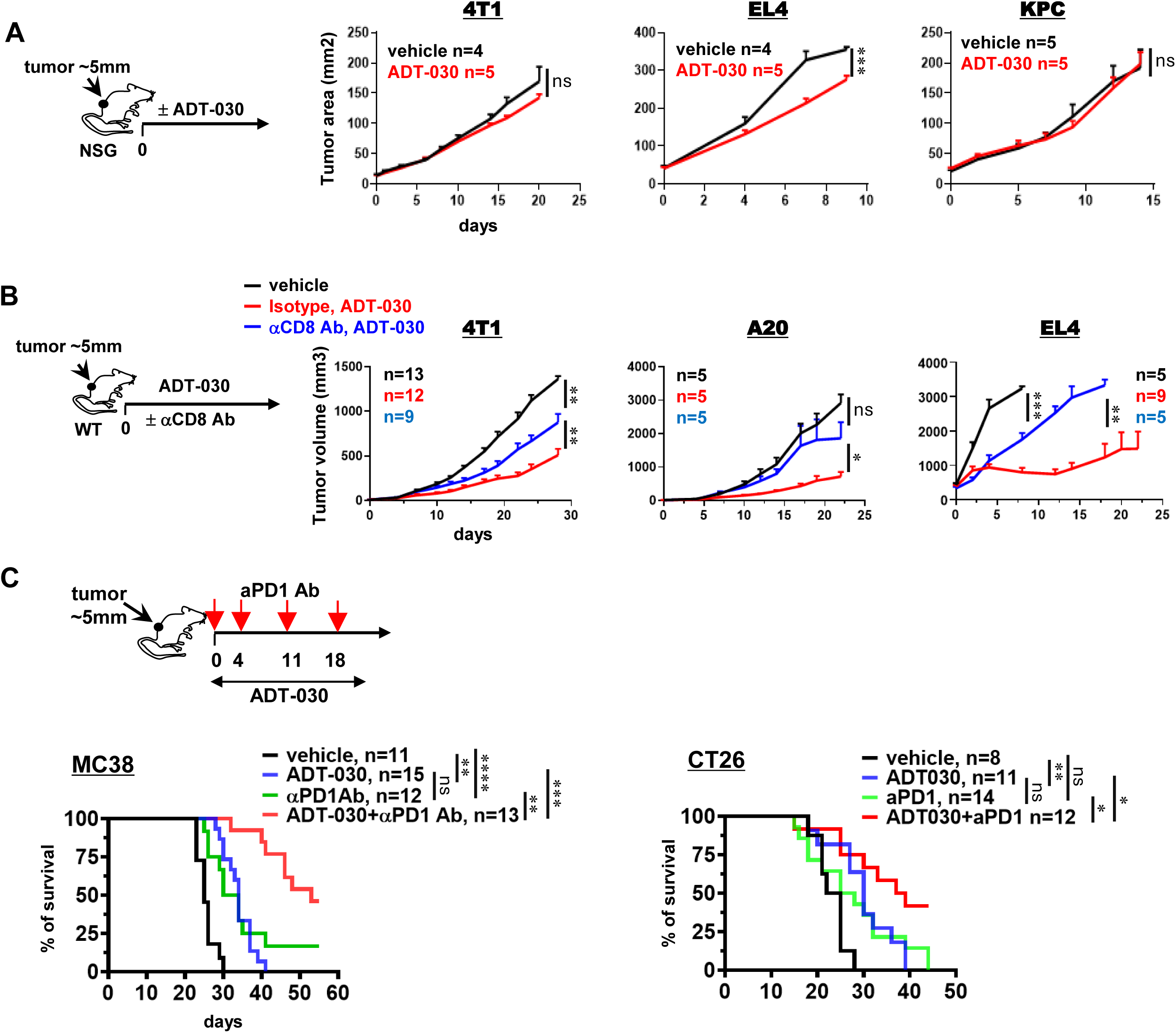
ADT-030–mediated tumor control is immune-dependent and is enhanced by anti–PD-1 therapy. (A) Tumor growth curves of vehicle- or ADT-030–treated tumor-bearing NSG mice. The schema depicts the experimental design. Data from representative tumor models are presented as mean ± SEM of tumor area over time, with the number of mice in each group indicated. (B) CD8 T cell depletion attenuates the antitumor effect of ADT-030 in immunocompetent mice. As depicted in the schema, tumor-bearing mice received isotype control or CD8-depleting Ab before and during ADT-030 treatment. Vehicle-treated tumor-bearing mice were included as controls. Tumor growth curves are shown for the indicated tumor models and presented as mean ± SEM with the number of mice indicated. (C) Combination treatment with ADT-030 and aPD-1 Ab produces enhanced antitumor activity in MC38 and CT26 tumor models. As depicted in the schema, mice bearing s.c. MC38 or CT26 tumors received the indicated treatments. aPD-1 Ab (200 μg/mouse) was administered i.p. for four doses, and ADT-030 was given daily by oral gavage for 21 days. Kaplan–Meier survival curves are shown with the number of mice per group indicated. ns, not significant; *, P<0.05; **, P<0.01; ***, *P* < 0.001; ****, P<0.0001.

### ADT-030 treatment increases tumor immunogenicity to facilitate immune activation

Certain chemotherapeutic agents can induce immunogenic cell death (ICD), a regulated form of tumor cell death characterized by the exposure or release of danger-associated molecular patterns (DAMPs) that promote dendritic cell (DC) recruitment, activation, and subsequent priming of tumor-specific CD8⁺ T cells^24,25^. Given our observation that the antitumor efficacy of ADT-030 is immune-dependent, we hypothesized that ADT-030 treatment enhances tumor immunogenicity by inducing ICD. We therefore investigated whether ADT-030 treatment elicits canonical hallmarks of ICD in tumor cells, including cell surface exposure of calreticulin (CRT), and extracellular release of ATP and high-mobility group box 1 (HMGB1). Flow cytometric analysis revealed that ADT-030 treatment significantly increased CRT exposure on the surface of dying tumor cells across multiple tumor cell lines compared with vehicle-treated controls (Fig. 6A and Supplemental Fig. 6). Consistently, luminescence-based assays detected elevated extracellular ATP levels (Fig. 6B), and ELISA confirmed enhanced HMGB1 release following ADT-030 treatment (Fig. 6C).

**Fig. 6.**
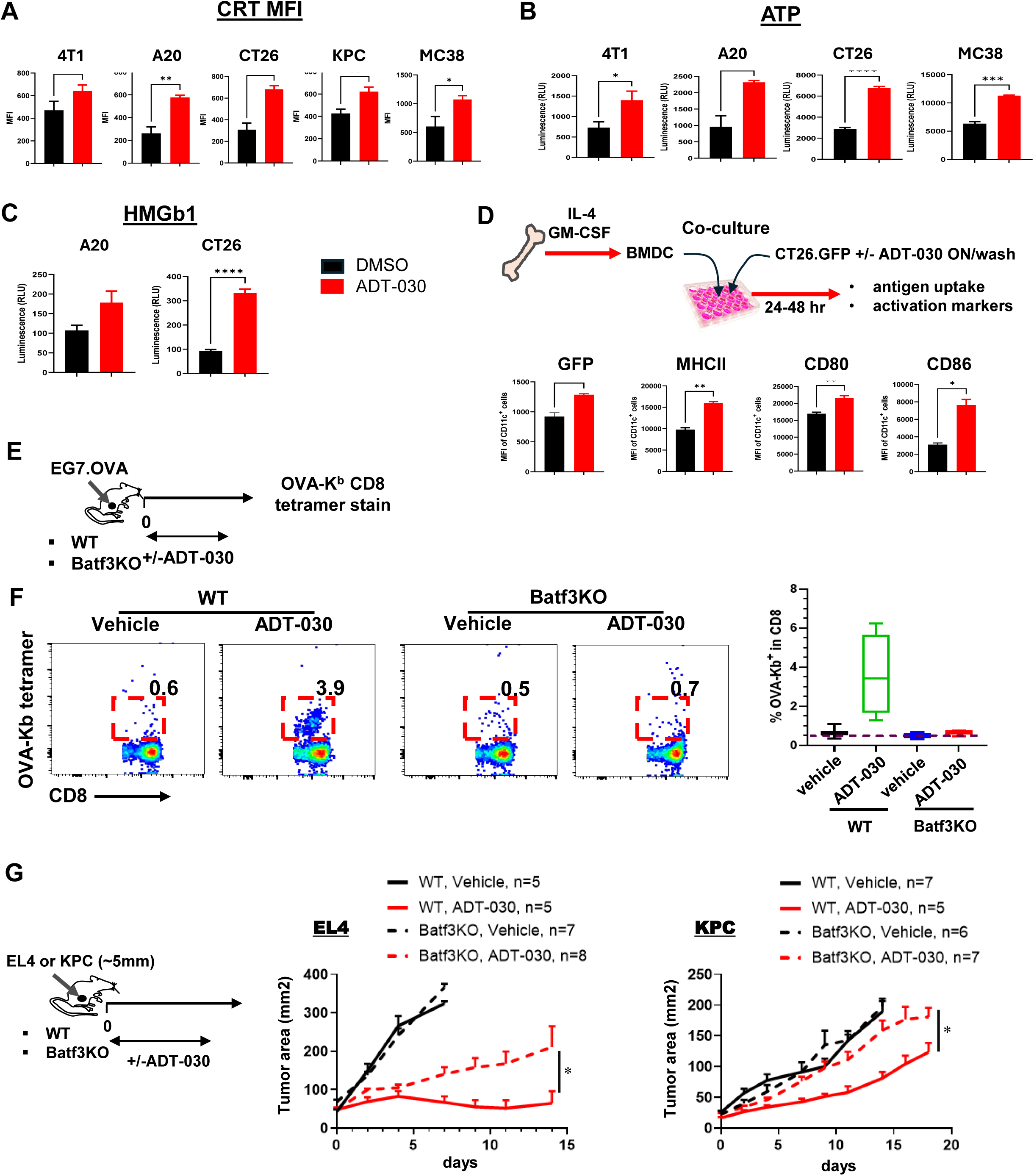
ADT-030 treatment enhances tumor immunogenicity by inducing immunogenic tumor cell death, promoting DC maturation, and augmenting CD8^+^ T cell priming. (A-C) Detection of ICD markers. Tumor cells were treated with DMSO or ADT-030 for 24 hours. Cell surface expression of CRT was detected by FACS, and MFI values are summarized in (A) as mean ± SEM. Extracellular ATP and HMGB1 levels from the indicated cells were measured and results shown as mean +/- SEM are summarized in (B) and (C), respectively. (D) ADT-030-treated tumor cells promote DC maturation. The schema depicts the experimental setup in which BMDCs were co-cultured with ADT-030 or DMSO pre-treated 4T1.GFP cells for 48 hrs. Uptake of the GFP signal by DCs and DC activation markers (MHC-II, CD80 and CD86) were assessed by FACS. MFIs are summarized as mean ± SEM in bar graphs. (E, F) cDC1-dependent priming of tumor-specific CD8⁺ T cells. The schema depicts the experimental procedures (E). EG7.OVA tumor cells were s.c. implanted to the flanks of WT or Batf3KO mice. Mice with palpable tumors were treated with vehicle or ADT-030. Peripheral blood was collected 10 days after treatment to detect OVA-specific CD8+ T cells by OVA-Kb tetramer. Representative dot plots are shown in (F). Numbers of the gated region demark percent of tetramer-positive cells in CD8+ T cell population. Results shown as mean +/- SEM are summarized in the bar graph. (G) ADT-030 efficacy in mice diminishes in the absence of cDC1. As depicted in the schema, EL4 or KPC cells were s.c. implanted to WT or Batf3KO mice. Mice with palpable tumors were treated daily with vehicle or ADT-030 daily till the endpoint. Tumor growth curves are shown as mean ± SEM, with the number of mice per group indicated. *, *P* < 0.05; **, *P* < 0.01; ***, *P* < 0.001; ****, *P* < 0.0001.

We next assessed whether these ICD-associated changes translated into enhanced DC activation. GFP-expressing CT26 cells were pretreated with vehicle or ADT-030 and subsequently co-cultured with bone marrow–derived DCs (BMDCs). BMDCs co-cultured with ADT-030-treated CT26.GFP cells displayed increased antigen uptake, as measured by GFP fluorescence, and exhibited a more mature and activated phenotype, characterized by elevated expression of MHC II, CD80, and CD86 (Fig. 6D), indicating that ADT-030-induced tumor cell death promotes DC activation.

To determine whether the antitumor efficacy of ADT-030 requires DC-dependent priming of tumor-specific CD8⁺ T cells in vivo, we treated wild-type (WT) or Batf3-deficient (Batf3KO) mice bearing EG7.OVA tumors (Fig. 6E). Batf3KO mice lack type 1 conventional dendritic cells (cDC1), a DC subset essential for cross-presentation of tumor antigens to CD8⁺ T cells^26^. This experimental design enabled direct assessment of tumor antigen–specific CD8⁺ T-cell priming by tracking OVA-specific CD8⁺ T cells using an OVA-H-2Kᵇ tetramer, providing a readout of efficient DC-mediated antigen capture and cross-priming in response to ADT-030 treatment. OVA-specific CD8⁺ T cells were sparse in the circulation of vehicle-treated mice but were readily detectable in ADT-030-treated WT mice (Fig. 6F, WT group). In contrast, ADT-030-induced expansion of OVA-specific CD8⁺ T cells was abolished in Batf3KO mice (Fig. 6F, Batf3KO group), demonstrating a reliance on cDC1 for ADT-030–mediated CD8⁺ T-cell priming. Consistent with the induction of tumor-specific CD8^+^ T cells, ADT-030 mediated robust tumor growth inhibition in WT mice, with 6 out of 8 mice achieving complete tumor regression. These mice were resistant to subsequent tumor rechallenge, indicating the establishment of protective immune memory (Supplemental Fig. 6A). In contrast, the efficacy of ADT-030 against EG7.OVA tumor was markedly diminished in Batf3KO mice (Supplemental Fig. 7B). Although EG7.OVA tumors are intrinsically more immunogenic than the parental EL4 line due to expression of a non-endogenous antigen, similar dependence on cDC1 was observed in additional tumor models. Specifically, ADT-030 efficacy was significantly reduced in Batf3KO mice bearing EL4 or KPC tumors (Fig. 6G), neither of which express exogenous model antigens, confirming that cDC1-dependent antitumor immunity is a general requirement for ADT-030 activity.

Collectively, these findings delineate the cellular cascade potentially initiated by ADT-030 treatment, encompassing induction of immunogenic tumor cell death, activation of cross-priming DCs, and subsequent expansion of tumor-reactive CD8⁺ T cells, thereby providing a mechanistic basis for the immune-dependent antitumor efficacy of ADT-030 in vivo.

### ADT-030 profoundly remodels the tumor immune microenvironment

Accumulating studies demonstrate that small-molecule inhibitors targeting oncogenic RAS signaling can remodel the tumor immune microenvironment (TiME) and elicit antitumor immune responses^14,27^. Given its RAS inhibition activity, we hypothesized that ADT-030 exerts similar beneficial immunomodulatory effects. To gain deeper insight into how ADT-030 shapes the immune landscape within the tumor, lymphoid organs, and metastatic niche, we performed mass cytometry (CyTOF) immune profiling of mammary tumors, spleens, and lungs collected from the orthotopic 4T1 breast cancer model (Fig. 7, schema). Tumor and spleen samples were collected at two time points (days 7 and 14) to monitor dynamic immunological changes during treatment, while lung samples were collected at day 14 to permit the development of metastatic disease within this timeframe. A 33-antibody panel was used to identify major immune populations, including CD8⁺ and CD4⁺ T cells, B cells, NK cells, DCs, neutrophils, monocytes, macrophages, and eosinophils (Supplemental Fig. 8A). After 7 days of treatment, ADT-030-treated tumors exhibited a marked reduction in myeloid populations, including neutrophils, monocytes, and eosinophils, compared with vehicle-treated controls; In contrast, the relative abundance of CD8⁺ T cells, CD4⁺ T cells, B cells, DCs, and macrophages increased in ADT-030-treated tumors (Fig. 7A-B). It is important to note that at this early time point, tumor sizes were comparable between treatment groups, minimizing potential confounding effects caused by differences in tumor burden. These immune alterations persisted after 14 days of treatment (Fig. 7B, d7 versus d14). Similar immune remodeling, including reduced neutrophil frequencies and increased proportions of CD8⁺ T cells, CD4⁺ T cells, and B cells, was observed in the spleens and lungs of ADT-030-treated mice (Supplemental Fig. 8B-E). Conventional flow cytometry analysis of day 7 tumor samples validated the CyTOF results, confirming reduced tumor-infiltrating neutrophils and monocytes along with increased CD8⁺ and CD4⁺ T cell frequencies (Fig. 7C). Moreover, ADT-030 enhanced the effector function of tumor-infiltrating CD8⁺ T cells, as evidenced by increased granzyme B and interferon-γ production (Fig. 7D). Multiplex immunofluorescence stain further confirmed the abundance of CD8+ T cells in the tumor at the expense of myeloid cells after ADT-030 treatment (Fig. 7E). Together, these data demonstrate that ADT-030 induces a durable and systemic remodeling of the TiME, shifting tumors toward a more immune “hot” phenotype.

**Fig. 7.**
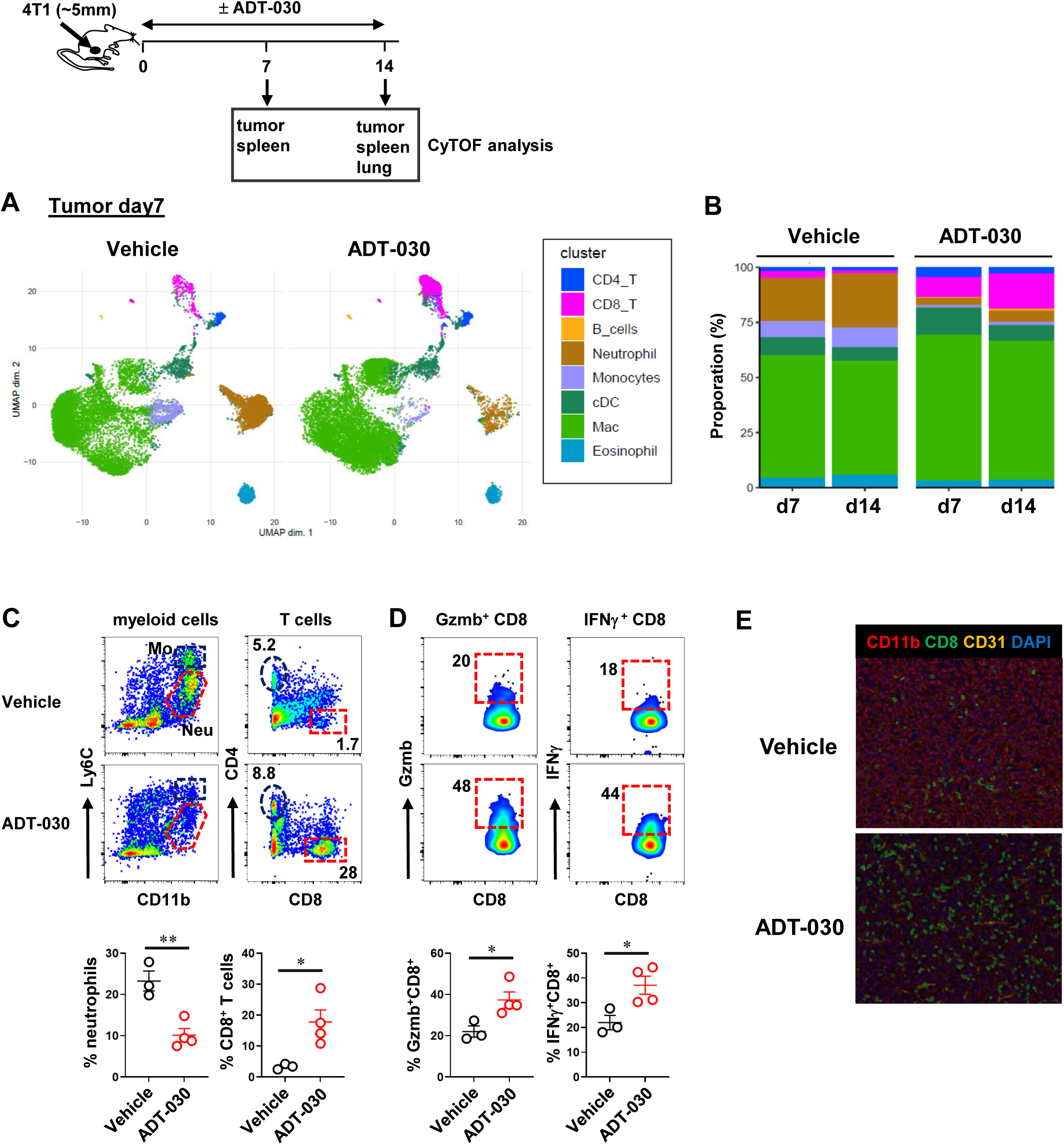
ADT-030 profoundly remodels the TIME. The schema depicts the experimental workflow for CyTOF-based immune profiling of tissue samples. (A) UMAP projection of CD45^+^ immune cells in tumors from mice treated with vehicle or ADT-030 for 7 days. Tumors collected after 14 days were analyzed in parallel. (B) Stacked bar plots showing the relative proportions of immune cell subsets among CD45⁺ cells in tumors at different time points following the indicated treatment. (C) Day 7 tumor samples analyzed by conventional flow cytometry to assess the abundance of myeloid cells, CD8⁺ T cells, and CD4⁺ T cells. Representative dot plots show the gating strategy, and frequencies of neutrophils and CD8⁺ T cells are summarized as mean ± SEM in bar graphs. (D) ICS of CD8⁺ T cells for GzmB and IFN-γ following PMA/ionomycin stimulation. Representative dot plots are shown, with quantification presented as mean ± SEM. *, *P* < 0.05; **, *P* < 0.01. (E) Representative immunofluorescence images of tumor sections stained with DAPI and antibodies against CD11b and CD8.

### ADT-030 normalizes the immune landscape at the metastatic site

To investigate whether ADT-030 can also reprogram TiME at metastatic sites, we performed single-cell RNA sequencing (scRNA-seq) on lung tissues from the 4T1 experimental metastasis model, with lungs from vehicle- or ADT-030-treated tumor-free mice included as controls (Fig. 8 schema). The t-SNE plot reveals distinct immune cell populations defined by their transcriptomic profiles (Fig. 8A and Supplemental Fig. 9). In tumor-bearing mice, ADT-030 treatment markedly reduced neutrophil abundance while increasing the frequencies of CD8⁺ T cells, CD4⁺ T cells, B cells, and NK cells within the metastatic lungs (Fig. 8B). In contrast, these compositional changes were marginal in the lungs of tumor-free mice (Fig. 8C). Overall, ADT-030 appeared to restore the immune cell landscape of tumor-bearing lungs toward a composition resembling that of tumor-free controls. Given the pronounced effect of ADT-030 on neutrophils, we next examined neutrophil-intrinsic transcriptional changes. Hallmark pathway analysis of paired samples revealed that ADT-030 altered the expression of 225 genes across 10 signaling pathways in neutrophils from tumor-bearing lungs, including type I and type II interferon responses, inflammatory signaling, p53 signaling, and apoptosis, whereas substantially fewer genes (65) and pathways (3) were affected in neutrophils from tumor-free lungs (Fig. 8D). Consistent with these findings, ADT-030 treatment upregulated interferon-α– and interferon-γ–responsive gene programs in neutrophils from tumor-bearing mice (Fig. 8E). Together, these results suggest that ADT-030 selectively reprograms immunosuppressive myeloid populations within metastatic lesions, contributing to normalization of the metastatic TiME while preserving immune architecture in non-tumor tissues.

**Fig. 8.**
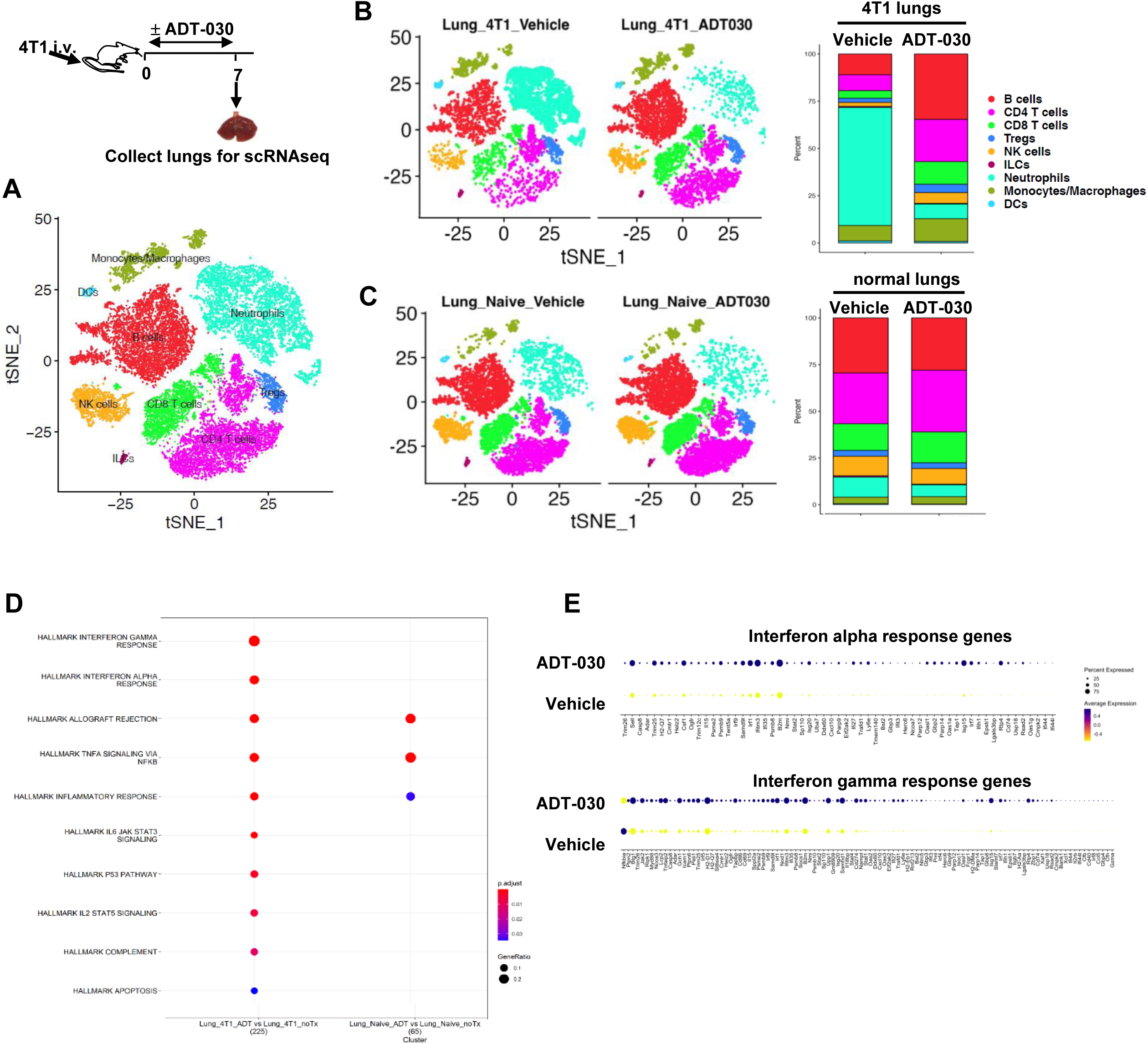
ADT-030 normalizes the immune landscape at the metastatic site. The schema depicts the workflow for scRNA-seq analysis of lung samples from the 4T1 experimental metastasis model. (A) t-SNE visualization of immune cell subsets among CD45⁺ cells in the lungs defined based on transcriptional profiles. (B) t-SNE plots showing annotated immune cell populations in lung samples across the indicated treatment conditions. The relative proportions of immune cell subsets among CD45⁺ cells are summarized in stacked bar plots. (C) Dot plots depicting signaling pathways in lung neutrophils that are differentially regulated by ADT-030 in tumor-bearing versus tumor-free mice. (D) Dot plots comparing the expression of interferon-α and interferon-γ response gene signatures in lung neutrophils from vehicle- and ADT-030-treated mice.

### ADT-030 selectively induces apoptosis in tumor-associated myeloid cells

Our immune profiling studies revealed that ADT-030 profoundly reduced neutrophils, the predominant myeloid subset in the 4T1 tumor model. This prompted us to investigate whether neutrophils in tumor-bearing mice are preferentially sensitive to ADT-030 compared with other immune populations. To address this, splenocytes from mice bearing well-established 4T1 tumors, containing both lymphoid cells (including CD8⁺ T cells) and myeloid cells (predominantly neutrophils), were treated in vitro with increasing concentrations of ADT-030 or DMSO control (Fig. 9A). Overnight exposure to ADT-030 induced robust apoptosis in myeloid cells at all doses tested, whereas CD8⁺ T cells were largely unaffected (Fig. 9B-C), indicating selective sensitivity of myeloid cells to ADT-030. To determine whether this selectivity can be recapitulated in vivo, we analyzed tail blood from 4T1-bearing mice treated with vehicle or ADT-030 for 7 days, with naïve (tumor-free) mice included as controls (Fig. 9D). ADT-030 treatment significantly increased the frequency of apoptotic myeloid cells in tumor-bearing mice compared with vehicle-treated tumor-bearing mice or naïve controls; In contrast, the frequency of apoptotic CD8⁺ T cells remained at baseline levels across all groups (Fig. 9E-F). Consistent with these findings, myeloid cells from ADT-030-treated tumor-bearing mice exhibited significantly reduced p-ERK and p-p38, along with increased p-p53 and cCas3 (Fig. 9G). These signaling changes were not observed in CD8⁺ T cells from the same animals (Fig. 9H). In the EL4 tumor model, which also features progressive accumulation of immunosuppressive myeloid cells^28^, ADT-030 similarly induced selective apoptosis within the myeloid cell compartment (data not shown). Collectively, these results demonstrate that ADT-030 selectively induces apoptosis in tumor-associated myeloid cells while sparing CD8⁺ T cells, providing a mechanistic explanation for its ability to remodel the TiME without compromising antitumor T cell immunity.

**Fig. 9.**
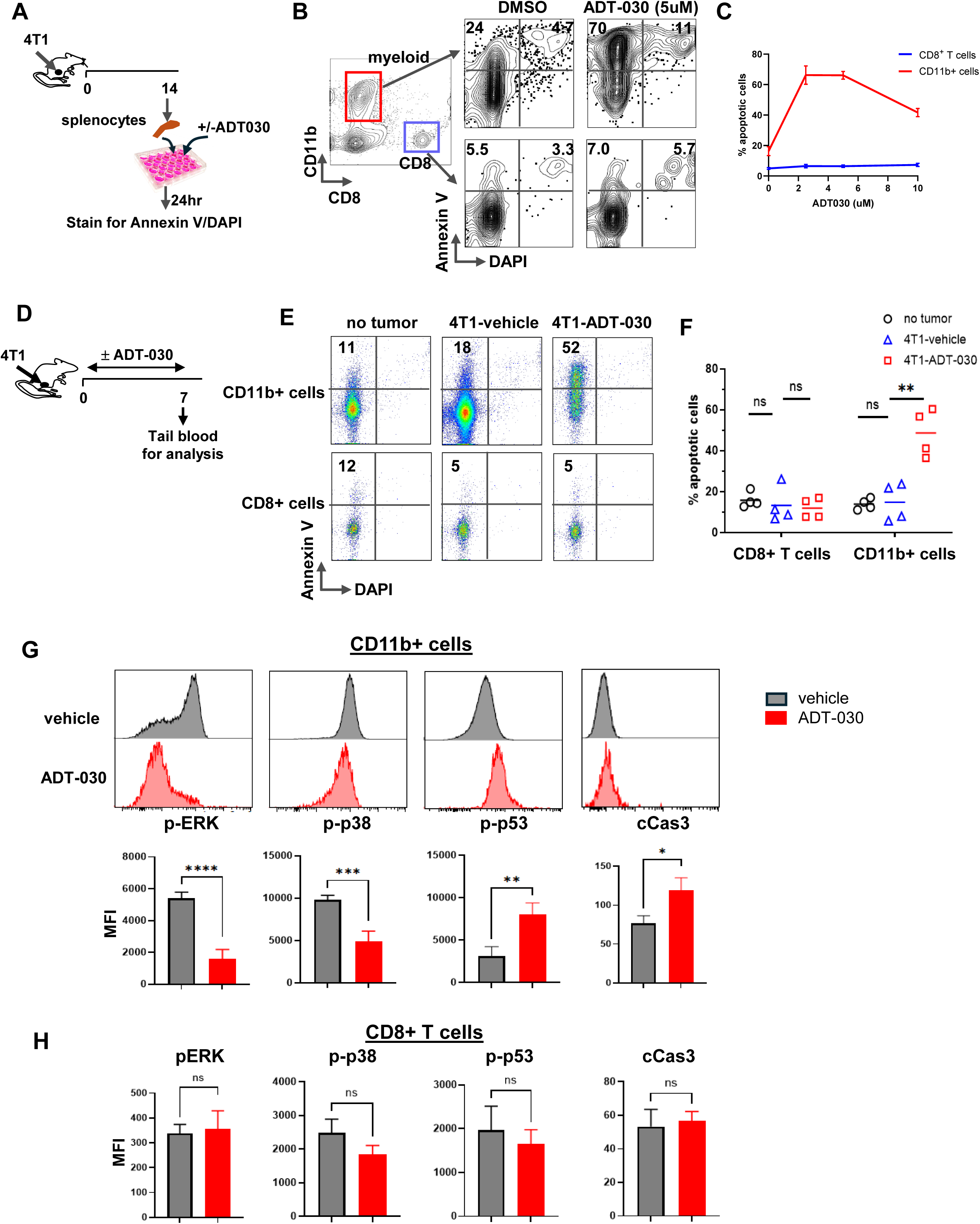
ADT-030 selectively induces apoptosis in tumor-associated neutrophils. (A) Schema depicting the workflow for assessing neutrophil sensitivity to ADT-030 ex vivo. Peripheral blood was collected via tail vein from mice bearing advanced 4T1 tumors (>10 mm). White blood cells were cultured and treated with DMSO or increasing concentrations of ADT-030 for 16 h, followed by assessment of apoptosis by FACS. (B) Representative dot plots showing apoptosis in myeloid cells and CD8⁺ T cells. Numbers indicate the percentage of apoptotic cells within the gated populations. (C) Quantification of apoptotic myeloid cells and CD8⁺ T cells under each culture condition, presented as mean ± SD (n = 6 replicates). (D) Schema depicting the in vivo assessment of apoptosis in circulating immune cells. Mice bearing established orthotopic 4T1 tumors were treated with vehicle or ADT-030 for 7 days. Peripheral blood was collected for apoptosis analysis by FACS. Blood from tumor-free naïve mice served as controls. (E) Representative dot plots showing apoptosis in circulating myeloid cells and CD8⁺ T cells. Numbers indicate the percentage of apoptotic cells. (F) Summary of apoptotic cell frequencies in myeloid cells and CD8⁺ T cells under the indicated treatment conditions. (G) Representative histograms showing expression of p-ERK, p-p38, p-p53, and cCas3 in circulating myeloid cells from vehicle- or ADT-030-treated mice. MFI values are summarized in bar graphs (mean ± SEM). (H) Bar graphs summarizing MFI values of the indicated markers in circulating CD8⁺ T cells. ns, not significant; . *, *P* < 0.05; **, *P* < 0.01; ***, *P* < 0.001; ****, *P* < 0.0001.

## Discussion

In recent years, the development of inhibitors targeting components of the RAS signaling pathway, including RAF, MEK, ERK, and mutant RAS isoforms, has advanced rapidly, yielding a growing pipeline of promising candidate compounds. Extensive preclinical studies have evaluated the efficacy and toxicity of these agents across diverse tumor models, and several have progressed into clinical trials. Among them, two KRAS^G12C^ inhibitors, sotorasib and adagrasib, have received FDA approval for the treatment of malignancies harboring KRAS^G12C^ mutations. However, a persistent challenge associated with these inhibitors is the rapid emergence of drug resistance driven by various tumor-intrinsic and tumor-extrinsic mechanisms^15,29,30^. To address this limitation, the development of broader-spectrum inhibitors, including pan-KRAS and pan-RAS inhibitors, is actively being pursued and has shown promise in overcoming resistance^15^.

Our previous work demonstrated that PDE10A inhibition suppresses RAS-MAPK signaling in addition to attenuating Wnt/β-catenin transcriptional activity in several human cancer cell lines^1,2,6,7^. In the current study, we provide evidence that ADT-030, a potent and selective PDE10A inhibitor, reduces multiple downstream effectors of RAS signaling, including RAF, ERK, and AKT, across a broad panel of murine tumor cell lines. While some of the cell lines examined, such as CT26, EL4, and KPC, carry oncogenic KRAS mutations, the other cell lines, including 4T1, A20, B16, and MC38, lack mutant RAS^21^, yet all cell lines exhibit measurable basal RAS activity. Further supporting ADT-030 as a functional pan-RAS inhibitor, our parallel work demonstrates that ADT-030 exhibits potent cytotoxicity against murine and human pancreatic tumor cell lines harboring distinct KRAS mutations and retains activity in tumor cells that have acquired resistance to the KRAS^G12C^ inhibitors sotorasib or adagrasib^19^.

One central finding of our study is that ADT-030 has the ability to elicit antitumor immune responses. We delineate the mechanistic basis of this effect, linking ADT-030 treatment to the induction of tumor cell ICD, enhanced dendritic cell maturation, and augmented priming of tumor-specific CD8⁺ T cells. We further show that activation of the host immune system in turn contributes to ADT-030 efficacy, as evidenced by the reduced antitumor activity observed in immunodeficient mice or following CD8⁺ T-cell depletion. It is worth noting that ADT-030 retains partial antitumor activity in certain immunodeficient settings, including EL4 tumors in NSG mice (Fig. 5A) and A20 tumors in Rag2KO mice (Supplemental Fig. 5), indicating a tumor-intrinsic component to its efficacy. Moreover, CD8⁺ T cell depletion in immunocompetent mice tended to result in a partial loss of antitumor activity, an effect that was particularly evident in the 4T1 and EL4 models (Fig. 5B). Together, these findings suggest that the extent to which ADT-030 efficacy depends on host immunity is cancer type-dependent, and that immune populations beyond CD8⁺ T cells, including CD4⁺ T cells, B cells, NK cells, and M1-like macrophages, may also contribute to the overall antitumor response. Consistent with this interpretation, increased intratumoral abundance of these immune subsets was detected by CyTOF and scRNA-seq analyses (Figs. 7 and 8).

Our study provides compelling evidence that pharmacological inhibition of PDE10A can profoundly remodel the TiME. A key immunomodulatory effect of ADT-030 is the selective depletion of tumor-associated myeloid cells, the so-called MDSCs, as observed in the 4T1 tumor model. It is well documented that during 4T1 tumor progression, the myeloid compartment becomes increasingly dominated by neutrophils^28^. These tumor-associated neutrophils (TANs), often classified as granulocytic myeloid-derived suppressor cells (gMDSCs), not only potently suppress tumor-reactive T cells, B cells, and NK cells but also condition distant organs to facilitate metastatic seeding^31^. Our data demonstrate that ADT-030-mediated MDSC depletion restores intratumoral T, B, and NK cell populations, enhances T cell effector function, and disrupts tumor colonization at metastatic sites. Importantly, ADT-030 induces only mild immune alterations in tumor-free mice (Fig. 8C), suggesting minimal impairment of normal myelopoiesis. Moreover, ADT-030-mediated MDSC reduction does not result in neutropenia in tumor-bearing mice; instead, it normalizes the TiME by reshaping the immune landscape toward that observed in tumor-free mice (Fig. 8B). Notably, the residual neutrophils in ADT-030-treated mice acquire a phenotype characterized by an interferon-stimulated gene (ISG) signature (Fig. 8E). This observation is consistent with recent studies reporting that ISG^high^ neutrophils expand within tumors following successful immunotherapy in mouse models, and that their emergence in cancer patients is associated with improved therapeutic outcomes^32,33^. It has been reported that p38-mediated inactivation of interferon signaling is critical for neutrophils to acquire a MDSC phenotype and immune suppression activity^34^. Consistent with this mechanism, our finding that ADT-030 reduces p38 activation in MDSCs implicates p38 as a downstream signaling node affected by PDE10A inhibition (Fig. 9G). Collectively, these findings indicate that PDE10A inhibition leads to preferential targeting of aberrantly differentiated MDSCs that expand during tumor progression while sparing normal homeostatic myelopoiesis and promoting the emergence of a subset of antitumor neutrophils.

Highly potent and selective inhibitors of PDE10A have been reported to induce cell cycle arrest and apoptosis of human cancer cell lines grown *in vitro* at concentrations that activate cGMP/PKG signaling^1,5^. Accumulating evidence indicates that PDE10A inhibitor-mediated PKG activation can result in the suppression of tumor-intrinsic β-catenin transcriptional activity and RAS-MAPK signaling^2,7^. Mechanistically, suppression of β-catenin transcriptional activity appears to result from PKG-mediated phosphorylation of β-catenin leading to its ubiquitination and proteasomal degradation^5,6^. A similar mechanism may underlie ADT-030-mediated inhibition of multiple signaling pathways, including RAS–MEK–ERK, AKT–mTOR–RPS6K, and p38 MAPK pathways. Supporting this hypothesis, cGMP-elevating agents have been reported to activate PKG, leading to disruption of RAS membrane localization and activation, as well as interference with receptor tyrosine kinase activity, resulting in attenuated MAPK signaling^35–38^. The exact mechanisms by which PDE10A inhibition leads to suppression of multiple MAPK signaling pathways in cancer cells and MDSCs await further investigation.

ADT-030 exhibits a favorable toxicity profile in preclinical models, distinguishing it from earlier PDE10A inhibitors developed for central nervous system (CNS) disorders such as Huntington’s disease and schizophrenia. We demonstrate in mouse tumor models that ADT-030 is orally bioavailable and accumulates in circulation, spleen, lung, and tumor at therapeutically relevant levels. Importantly, ADT-030 is barely detectable in the brain, where PDE10A is highly expressed^22^, suggesting a low risk of CNS toxicity. Mice treated with ADT-030 for up to 40 days exhibited no signs of toxicity, including weight loss, diarrhea, or behavioral changes. At the cellular level, ADT-030’s cytotoxic effects appear to be restricted to MDSCs, with minimal impact on CD8^+^ T cells. The mechanistic basis underlying the differential sensitivity of MDSCs versus CD8⁺ T cells to ADT-030 remains unclear but may relate to the fact that ADT-030 does not directly inhibit MAPK signaling. Instead, its regulatory effects are mediated through modulation of the cGMP-PKG axis resulting from PDE10A inhibition. We speculate that MDSCs require stringent regulation of intracellular cGMP levels and PKG activity to sustain the elevated MAPK signaling necessary for their differentiation, expansion, and survival. In contrast, CD8⁺ T cells may be less sensitive to fluctuations in cGMP and PKG signaling. Consequently, disruption of the cGMP-PKG axis by PDE10A inhibition may be more detrimental to MDSCs than to CD8⁺ T cells. Supporting this hypothesis, prior studies have shown that inhibition of PDE5, another cGMP-regulating phosphodiesterase, reduces MDSCs and their suppressive function^39^. Further studies are needed to elucidate the interplay among PDE10A activity, cGMP levels, PKG activation, and MAPK signaling in MDSCs compared with normal neutrophils and CD8⁺ T cells.

In conclusion, our study establishes PDE10A as a valid therapeutic target. By attenuating aberrant MAPK signaling within the tumor microenvironment, PDE10A inhibition holds the promise to simultaneously suppress cancer progression and unleash antitumor immunity. The development of potent and selective PDE10A inhibitors therefore represents a novel, safe and effective strategy to enhance cancer immunotherapy.

## Materials and Methods

### Cell lines and cell culture conditions

Murine tumor cell lines 4T1, A20, B16F10, CT26, EL4, EG7.OVA and MC38 were purchased from ATCC. The KPC pancreatic cancer cell line carrying mutations in p53 and Kras (G12D) was a gift from Dr. Lei Zheng (Johns Hopkins University). Luciferase-expressing 4T1 cells (4T1.luc) and GFP-expressing CT26 cells (CT26.GFP) were generated by transducing tumor cells with retroviruses carrying luciferase or GFP. All tumor cell lines were cultured in RPMI 1640 or DMEM (HyClone Laboratories) culture medium supplemented with 10% fetal bovine serum albumin (FBS), 1% penicillin/streptomycin (HyClone Laboratories), 1% non-essential amino acids, 1% glutamine and 0.1mM 2-mercaptoethanol at 37°C in a 5% CO2 incubator.

### Mice

BALB/c and C57/BL6 mice of 6–8 weeks old were purchased from the Charles River Laboratories. Batf3 knockout (Batf3KO) mice and NOD-Scid IL-2Rγ-null (NSG) mice were purchased from Jackson Laboratory. Rag2 knockout (Rag2KO) mice were purchased from Taconic Biosciences. All mice were housed under specific pathogen-free conditions by Laboratory Animal Services of Augusta University. All animal experiments and procedures were performed in accordance with the institutional protocol and were approved by the Institutional Animal Care and Use Committee (IACUC) of Augusta University.

### Antibodies and reagents

The following fluorochrome-conjugated antibodies were used for flow cytometry analysis: CD4 (GK1.5), CD8 (53-6.7), CD19 (6D5), PD1 (RMP1-14), CD11b (M1/70), I-A/I-E (M5/114.15.2), CD80 (16-10A1), CD86 (GL-1), Annexin V, PE-conjugated Donkey anti-rabbit IgG, anti–TNF-α (MP6-XT22), anti–IFN-γ (XMG1.2), anti–GM-CSF (MP1-22E9), anti-GzmB (GB11), anti-MEK1 Phospho (Ser298), anti-ERK1/2 Phospho (Thr202/Tyr204), anti-AKT Phospho (Ser473), anti-S6 Phospho (Ser235/Ser236), and anti-p38 MAPK Phospho (Thr180/Tyr182). Antibodies were purchased from Biolegend unless otherwise noted. Anti-phospho-p53 (Ser15) (D4S1H) antibody, anti-cleaved Caspase-3 (Asp175) (5A1E) antibody and anti-non-phospho (active) β-Catenin (Ser33/37/Thr41) (D13A1) antibody were purchased from Cellsignaling Technology. PE Anti-Calreticulin (CRT) antibody (EPR3924) was purchased from Abcam. Murine Gmcsf and IL4 were purchased from PeproTech. Anti-mouse PD1 antibody (RMP1-14) and anti-mouse CD8α (2.43) antibody were purchased from BioXcell.

The synthesis of ADT-030 (Z)-2-(5-methoxy-2-methyl-1-(3,4,5-trimethoxybenzylidene)-1H-inden-3-yl)-N-(1-methylpyrrolidin-3-yl) acetamide is based on a procedure originally described in U.S. patent 20200223815 using 3-(4-methoxyphenyl)-2-methylacrylic acid as the starting material.

### Cell preparation and flow cytometry analysis

Peripheral blood was collected via tail vein bleeding. Spleen and lymph node samples were processed into single-cell suspensions. Tumor and lung samples were dissociated into cell suspensions using the gentleMACS Octo Dissociator (Miltenyi Biotec) following the manufacturer’s instruction. Red blood cells were lysed by ACK lysing buffer (Quality Biological). For cell surface molecule detection, cells were stained with fluorochrome-conjugated antibodies for 15 minutes at room temperature in the dark. For detection of apoptosis, cells were harvested for Annexin V staining (BioLegend) following manufacturer’s instruction. 4′,6-diamidino-2-phenylindole (DAPI) was added to cells at 0.5µg/ml before flow cytometry analysis. For detection of surface calreticulin (CRT), tumor cells were treated with DMSO or 10uM ADT-030 at 37°C for 24 h. Cells were harvested and fixed at 4°C with 0.25% paraformaldehyde in PBS, washed, and stained with PE-conjugated anti-CRT Ab for 30 min at 4°C. Cells were washed and resuspended in PBS buffer with addition of DAPI before FACS analysis. For cytokine intracellular staining, T cells in splenocytes or tumor tissue suspensions were stimulated with the Leukocyte Activation Cocktail (BD Biosciences) for 3-4 hours at 37°C. Cells were harvested and surface stained, followed by cytokine staining using the Fixation/Permeabilization Solution Kit (BD Biosciences). For detection of cleaved caspase 3 and phosphorylated molecules, including AKT, S6, p38 and p53, cells were stained with the specified antibody following fixation and permeabilization with 4% formaldehyde and ice-cold methanol. FACS data were acquired using an Attune NxT Flow Cytometer (ThermoFisher Scientific) or an Acea NovoCyte Quanteon (Agilent). Flow data were analyzed using FlowJo software (Tree Star).

### Cell viability assay

Cancer cell viability was measured by PrestoBlue HS Cell Viability Reagent (Invitrogen) following manufacturer’s instruction. Briefly, cells were seeded in flat-bottom 96-well plate at the density of 10,000-20,000 cells in 100ul volume per well. After overnight culture, increasing doses of ADT-030 were added to specified wells with triplicate wells for each condition. PrestoBlue HS reagent was added to wells 24 hours later. Cell viability was evaluated by measuring the fluorescence using a BioTek Microplate Reader.

### Western blot analysis

Cells were lysed in ice-cold RIPA buffer supplemented with protease inhibitors (Thermo Scientific Chemicals). Protein concentrations were determined using the BCA assay kit (Pierce), and 20 µg of total protein per sample was resolved on a 10% SDS–PAGE gel (Bio-Rad) along with a molecular weight marker. Proteins were transferred onto a PVDF membrane using a wet transfer system at 200 mA for 2 hours. Membranes were blocked in 5% non-fat milk in TBST for 1 hour at room temperature and incubated overnight at 4 °C with the appropriate primary antibodies. After washing with TBST, membranes were incubated with HRP-conjugated secondary antibodies for 1 hour at room temperature. Protein bands were visualized using SuperSignal West Pico PLUS Chemiluminescent Substrate (Thermo Scientific) and imaged on a ChemiDoc system with automatic exposure settings. Densitometric analysis was performed using ImageJ, and protein levels were normalized to β-actin.

### RAS Pull-down assay

Detection of activated (GTP-bound) RAS protein in cell lysates was performed using an affinity pull-down approach based on the RAS-binding domain (RBD) of the RAS effector kinase RAF1. Tumor cells grown to 80-90% confluency were lysed in ice-cold lysis buffer. Lysates were collected after centrifugation at 13,500g for 5 minutes at 4 °C. For RAS-GTP measurement, approximately 500 µg of total protein from each sample was incubated with the Raf-RBD beads (Cytoskeleton Inc., catalog #RF-02). The assay was performed according to the manufacturer’s instructions, with minor modifications to the lysis buffer (50 mM Tris-HCl, pH 7.5; 150 mM NaCl; 1% NP-40; 10% glycerol) supplemented with protease and phosphatase inhibitors. Following incubation and washing, bead-bound RAS-GTP was detected by Western blotting using pan-RAS antibodies (Proteintech, catalog #60309-1-Ig). Densitometric analysis was performed using ImageJ to determine the ratio of RAS-GTP to total RAS across at least 3 biological replicates.

### Tumor cell extracellular ATP release assay

Tumor cells were seeded into a 96-well flat-bottom plate at a density of 15,000 to 20,000 cells per well in complete media. The cells were treated with DMSO or 10 μM ADT-030 in the presence of the RealTime-Glo™ Extracellular ATP Assay Reagent (Promega). Luminescence signal reflecting the extracellular ATP level was measured by SpectraMax (Molecular Devices).

### HMGB1 detection by ELISA

Tumor cells were treated with DMSO or 10 μM ADT030 at 37°C for 24 or 48 hours. Cell culture supernatants were collected for HMGB1 measurement by Lumit™ HMGB1 (Human/Mouse) Immunoassay kit (Promega) following the manufacturer’s instruction.

### Measurements of tumor antigen uptake by bone marrow-derived dendritic cells (BMDCs) and DC maturation markers

Bone marrow cells were harvested from the femurs and tibias of naïve BALB/c or C57BL/6 mice. After removal of red blood cells, bone marrow cells were cultured in complete media containing 25 ng/mL murine Gmcsf and 20 ng/mL murine IL-4 in a 6-well plate at 37°C and 5% CO_2_. Culture medium was replaced every 3 days. BMDCs were harvested on day 7-9 and used for subsequent experiments. GFP-expressing tumor cells, were treated with DMSO or ADT030 (10uM) overnight. Tumor cells were washed thoroughly to remove residual chemicals and then co-cultured with BMDCs at 2:1 ratio for an additional 24-48 hours. Cells were harvested and stained with antibodies against CD11c, MHCII, CD80 and CD86 for flow cytometry analysis. The uptake of tumor-derived antigens by BMDCs was evaluated by acquisition of GFP signal in CD11c+ cells. The maturation status of BMDCs was evaluated by the levels of MHCII, CD80 and CD86 in CD11c+ cells.

### Animal tumor models and in vivo treatments

Tumor cells were subcutaneously implanted to the right flank of mice. For B16F10, CT26, KPC and MC38 tumors, 0.1 x 10^6^ – 0.5 x 10^6^ tumor cells in 50 µl PBS were injected per mouse. For A20, EL4 and EG7.OVA tumors, 1.0 x 10^6^ – 2.0 x 10^6^ tumor cells were injected for each mouse. For orthotopic 4T1 tumor model, 0.1 x 10^6^ 4T1 cells in 50 µl PBS were implanted to the 3^rd^ mammary fat pad of female BALB/c mice. When tumor sizes became palpable (3-5 mm), mice were randomly assigned into groups to receive the specified treatments. The tumor size was monitored by caliper measurement every other day and expressed as the product of two perpendicular diameters in square millimeters. For 4T1 experimental metastasis model, 0.1 x 10^6^ 4T1.luc cells were injected to female BALB/c mice via tail vein.

ADT-030 was prepared in Maalox suspensions and given to mice daily via gavage at the dose of 150 mg/kg. For depletion of endogenous CD8^+^ T cells, anti-CD8α antibody (100 µg per injection/mouse) was injected intraperitoneally (i.p) to mice once a week for 4 weeks, with the first injection one day before ADT-030 treatment. For anti-PD1 antibody treatment, anti-PD1 antibody was injected into mice via i.p., 100 µg per injection, at the specified times.

### *In vivo* and *ex vivo* bioluminescence imaging (BLI)

BLI was performed for mice receiving i.v. injection of 4T1.luc tumor cells to assess tumor burden. Briefly, each mouse received i.p. injection of luciferin at 150 mg/kg and was anesthetized by inhalation of 2% isoflurane. Pseudocolor luminescent images were acquired and overlaid with photographic images using the Ami X system (Spectral Instruments Imaging). In some experiments, mice were euthanized after live imaging, and lungs from these mice were immediately collected into a 12-well plate and immersed in luciferin-containing PBS for ex vivo imaging. All BLI data were analyzed using the Aura Imaging Software (Spectral Instruments Imaging). The luminescence signal was quantified as photon/sec as an indicator of the tumor density in the lungs of mice.

### Mass Cytometry (CyTOF) procedure and data analysis

Spleen, tumor and lung samples were processed as described above. Cells were washed in Maxpar PBS and stained with Maxpar OnDemand Mouse Immune Profiling Panel Kit (Standard Biotools Part#920001) following the manufacturer’s protocol. Briefly, cells were first stained with 103Rh at a final concentration of 1μM in Maxpar PBS. Prior to surface staining, anti-mouse CD16/32 antibody (Biolegend Cat#101302) was added to cell suspension in ice-cold Maxpar cell staining buffer to block the Fc receptors for 10 minutes. Samples were stained with different single isotopic tagged (89Y-, 106Cd-, 110Cd-, 112Cd-) anti-mouse CD45 in Maxpar cell staining buffer for 30 minutes on ice followed by two washes. The surface antibody cocktail was then added into the pooled sample for 30 minutes. The cells were then washed and fixed with 2% paraformaldehyde for 15min at room temperature. After fixation, cells were washed in staining buffer, permeabilized and stained with DNA intercalator containing natural abundance Iridium (191/193Ir) prepared to a final concentration of 125nM in Maxpar Fix and Perm Buffer. Cells were washed in staining buffer, with subsequent washes in Maxpar Cell Acquisition Solution Plus (CAS+). Cells were resuspended in CAS+, filtered through a 35 μm nylon mesh filter cap and supplemented with a 1:10 dilution of EQ Six Element Calibration beads (Standard Biotools Part#201245). Samples were acquired on a CyTOF XT Mass Cytometer (Standard Biotools) at an event rate of 500 events/second or less. Mass cytometry data files were bead-normalized in CyTOF software v9.0 and processed data were cleaned up by gating live singlets and debarcoded by gating single CD45+ cells in FlowJo (v10) before algorithmic analysis. The CATALYST R package was employed to analyze the CyTOF data. Data normalization involved an arcsinh transformation with a cofactor of 5. Major cell types were identified from CD45^+^ live cells using the FlowSOM clustering method with default settings. These cell types were assigned based on lineage markers. To determine the optimal number of clusters, consensus clustering was performed for k values ranging from 2 to 30, guided by the relative decrease in the area under the CDF curve. Heatmaps depicting median protein expression and UMAP plots projecting the cell types were generated using the CATALYST package. Cell type abundances for each sample were calculated and visualized with bar or box plots.

### Bulk RNA-seq analysis

4T1 cells were treated with DMSO or 10 μM ADT-030 for 16 hours. Total RNA was extracted using a standard RNA isolation kit according to the manufacturer’s instructions. RNA quality was assessed using Bioanalyzer, and only samples with high RNA integrity were used for library preparation. RNA-seq libraries were generated using a poly(A) selection method and sequenced on an Illumina platform to obtain paired-end reads. Raw sequencing reads were aligned to the mouse reference genome (mm10) using a standard alignment pipeline. Gene-level read counts were generated and normalized across samples. Differential gene expression analysis was performed using a standard statistical framework, comparing ADT-030-treated samples to DMSO controls. Genes with adjusted p values < 0.05 were considered significantly differentially expressed. Pathway enrichment analysis was performed using the Reactome database. Separate analyses were conducted for upregulated and downregulated gene sets. Enriched pathways were ranked based on adjusted p-values and gene ratios.

### Single-cell RNA sequencing sample preparation, sequencing and data analysis

Lung tissues were processed into single cell suspensions as described above. Live immune cells were enriched by sorting CD45.2-positive cells using a Bigfoot Cell Sorter (ThermoFisher). Sorted cells were processed for scRNAseq libraries using the Chromium Controller (10X Genomics). scRNAseq libraries were generated for approximately 2,000 cells per sample using 10X 3’ single cell mRNAseq V3 reagents. Sequence was performed with Novaseq6000 System Illumina (Illumina Inc. San Diego, CA) platform following 10X Genomics guidelines. Each cell was tagged with a 16bp barcode sequence, which represents the identity of each single cell throughput of the analysis pipeline. The raw reads in the fastq format were processed using the 10X genomics cellranger analysis package. ADT-030 and vehicle samples were pooled using the cellranger aggr pipeline. The cellranger pipeline outputs containing gene-by-cell expression data from the aggregated libraries were imported into R package Seurat 3.1.4 to create a Seurat object(<u>86</u>). Quality control measures were implemented in Seurat to filter out cells expressing > 6000 genes.

Cells with a higher percentage of mitochondrial genes (percent of mt >0.3) were also excluded from the subsequent analysis. Normalization and scaling were performed within Seurat before subsequent analysis such as PCA, umap, tSNE, and clustering analyses. Visualization of the scRNA data was performed using Seurat tSNEplot, Vinplot, Featureplot, Dotplot, and DoHeatmap functions.

### Quantification of ADT-030 and PGE_2_ by liquid chromatography coupled with mass spectrometry (LC-MS)

For ADT-030 detection, tissue samples were homogenized in pre-chilled 80% acetonitrile together with 0.2ml stainless steel beads (0.1-0.3mm^3^), by high-speed beating using a Blue Bullet spin homogenizer with a speed setting 6 for 3 minutes in cold room (4°C). The homogenized sample was then centrifuged at 16,000g for 10 minutes at 4°C. For serum samples, pre-chilled acetonitrile was added to serum sample (final acetonitrile concentration is 80%), and then vortexed vigorously for 3 minutes at room temperature, followed by centrifugation at 16,000g for 10 min at 4°C. The supernatant was transferred to a new tube and centrifuged under vacuum till dry. The extracted samples were redissolved with 50% methanol for LC-MS analysis. Separation of ADT-30 was performed with a Phenomenex Kinetex C18 column (100x2.1mm, 1.7um) on a Shimadzu Nexera UHPLC system at a flowrate of 0.16ml/min, with a gradient elution from 20% to 98% acetonitrile (with 0.1% formic acid) in 5 minutes. The total analysis time is 12 minutes. The effluent was ionized via ion electrospray in positive mode on a TSQ Quantiva triple-quadrupole mass spectrometry with the following instrument settings: ion spray voltage 3500V, sheath gas 30, ion transfer tube temperature 325°C, aux gas 8, vaporizer temperature 250°C, and FWHM of 0.7 for both Q1/Q3 resolution. The optimal collision energy and RF lens were determined using commercial standards. Transitions 479.2/379.0 and 479.2/306.1 were used as qualifier and quantifier for ADT-030. The integrated peak areas for these transitions were calculated for each sample using Skyline software (version 20.0, University of Washington). PGE_2_ detection by LC-MS was similarly conducted using methanol for PGE_2_ extraction. Transitions 351.2/271.2 and 351.2/333.1 were used as qualifier and quantifier for PGE_2_.

### Statistics and reproducibility

Data were analyzed using Prism 10 (GraphPad Software Inc.). Experimental values are expressed as Standard Error Mean (±SEM). The statistical significance between two groups were determined using unpaired two-tailed Student’s *t* test. The statistical differences among three or more groups were determined using one-way analysis of variance (ANOVA) followed by Tukey’s multiple-comparison test. Statistical analysis for survival was determined by two sided Kaplan-Meier estimator with Mantel-Cox log-rank to compare curves using Prism. P values less than 0.05 were considered statistically significant.

## ACKNOWLEDGMENTS

We thank the Flow and Mass Cytometry Core Facility at Georgia Cancer Center (RRID: SCR_025747) for maintaining and ensuring the smooth operation of the flow cytometers. We acknowledge the support and contribution of the Integrated Genomics Core Shared Resources (RRID: SCR_026483) at the Georgia Cancer Center of Augusta University. We thank Dr. Wenbo Zhi at the Proteomics Core of Augusta University for technical assistance and support.

## Author Contributions

M.Y. G., Y.Y., O.D.O., X.W. S.E.S. and C.B. performed research and analyzed data; N.S. and A.A. contributed to data analysis; A.W.T.C., P.S., D.L., D.W., C.C.H. and H.S. analyzed data and edited the manuscript. G.P. and G.Z. conceived the study, designed research, analyzed data and wrote the manuscript.

---

**Supplemental Fig. 1.**
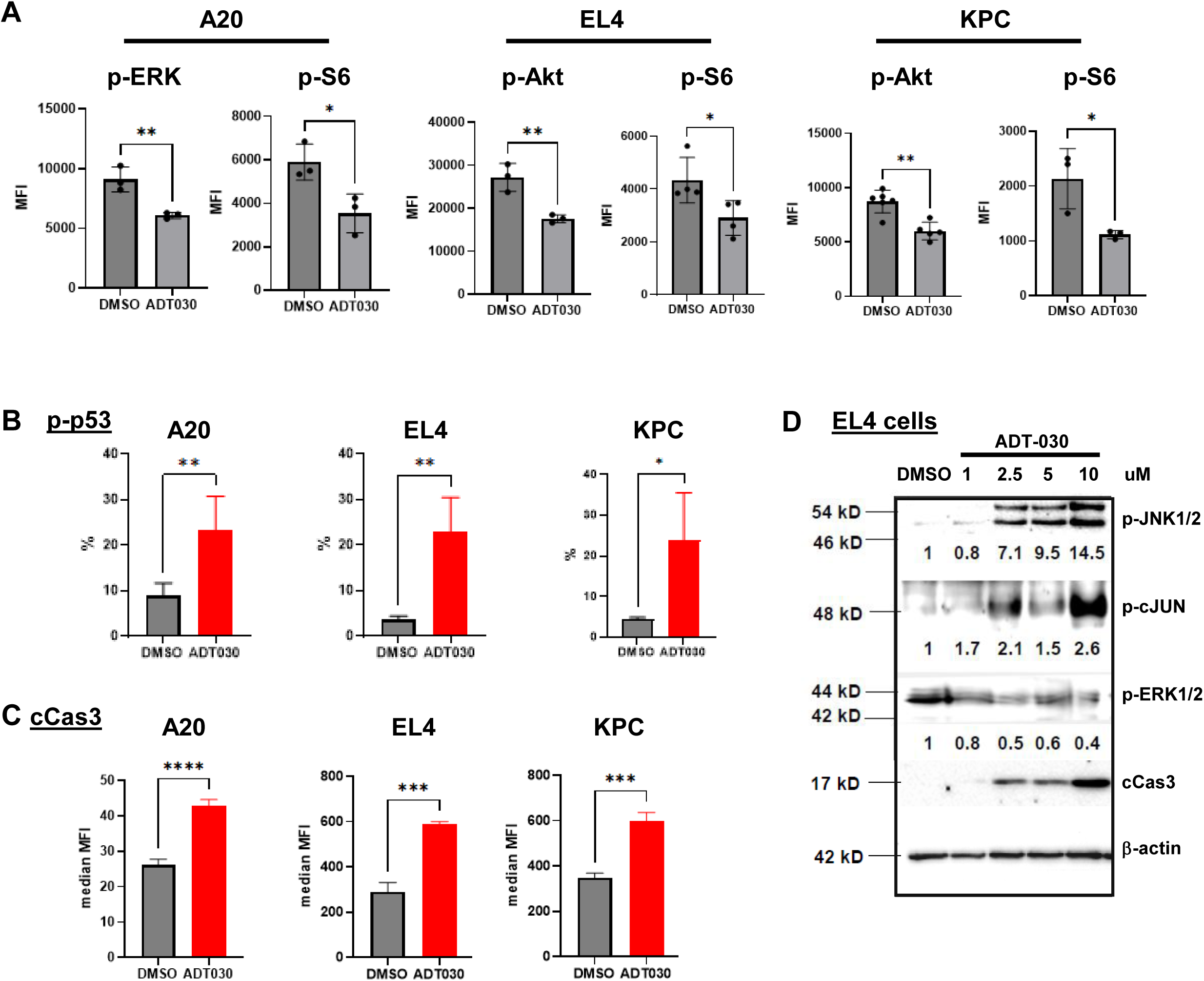
ADT-030 alters MAPK signaling and cell death pathways in multiple tumor cell lines. The indicated tumor cells were treated with either DMSO or ADT-030 for 16 hours prior to intracellular staining of the indicated molecules. (A) Summary of MFI of p-ERK, p-Akt and p-S6 in representative tumor cell lines. (B) Percent of phosphorylated p53 in representative tumor cell lines. (C) Summary of MFI of cCas3 in representative tumor cell lines. (D) WB analysis of phosphorylated JNK, c-JUN, ERK1/2, and cleaved caspase-3 in EL4 cells treated with escalating dose of ADT-030. β-actin serves as a loading control. *, *P* < 0.05; **, *P* < 0.01; ***, *P* < 0.001, ****, *P* < 0.0001.

**Supplemental Fig. 2.**
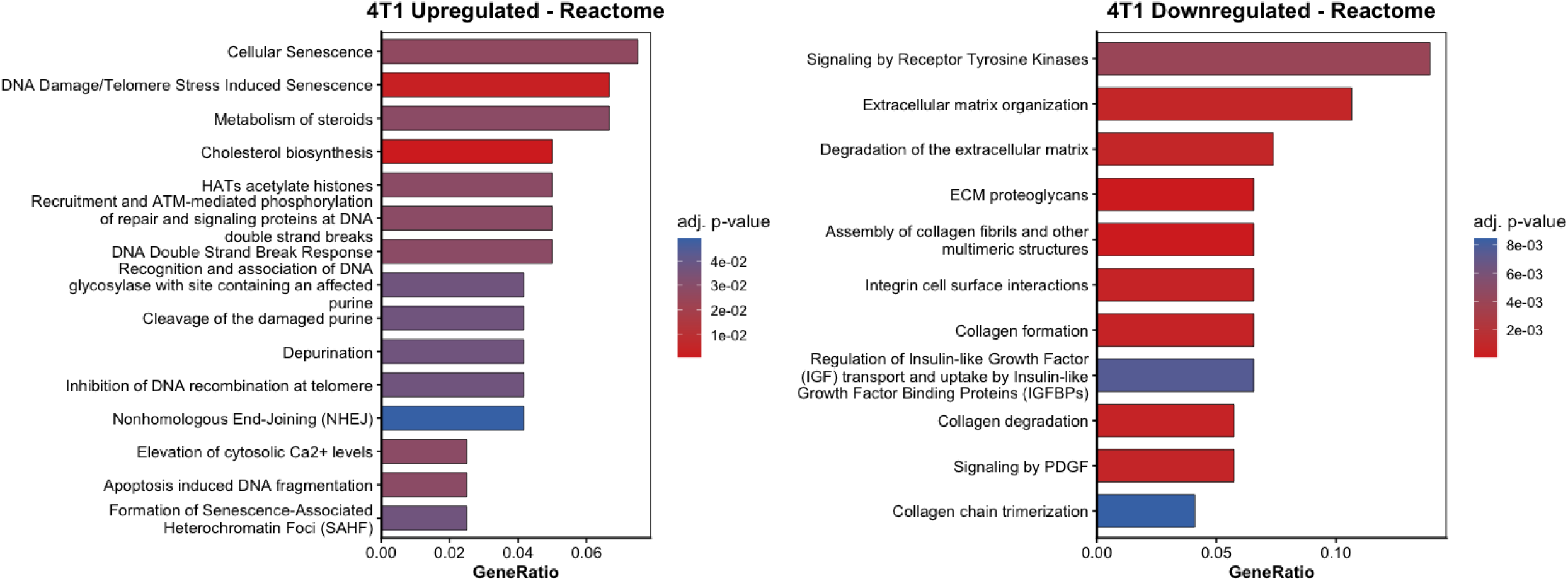
Pathways in 4T1 cells most affected by ADT-030 treatment. Pathway enrichment analysis was performed for 4T1 bulk RNA-seq. Pathway enrichment analysis was performed on differentially expressed genes from 4T1 cells treated with ADT-030 or DMSO. Bar plots show the most significantly upregulated and downregulated pathways. Gene ratio indicates the proportion of genes in each pathway. Color indicates adjusted p-values.

**Supplemental Fig. 3.**
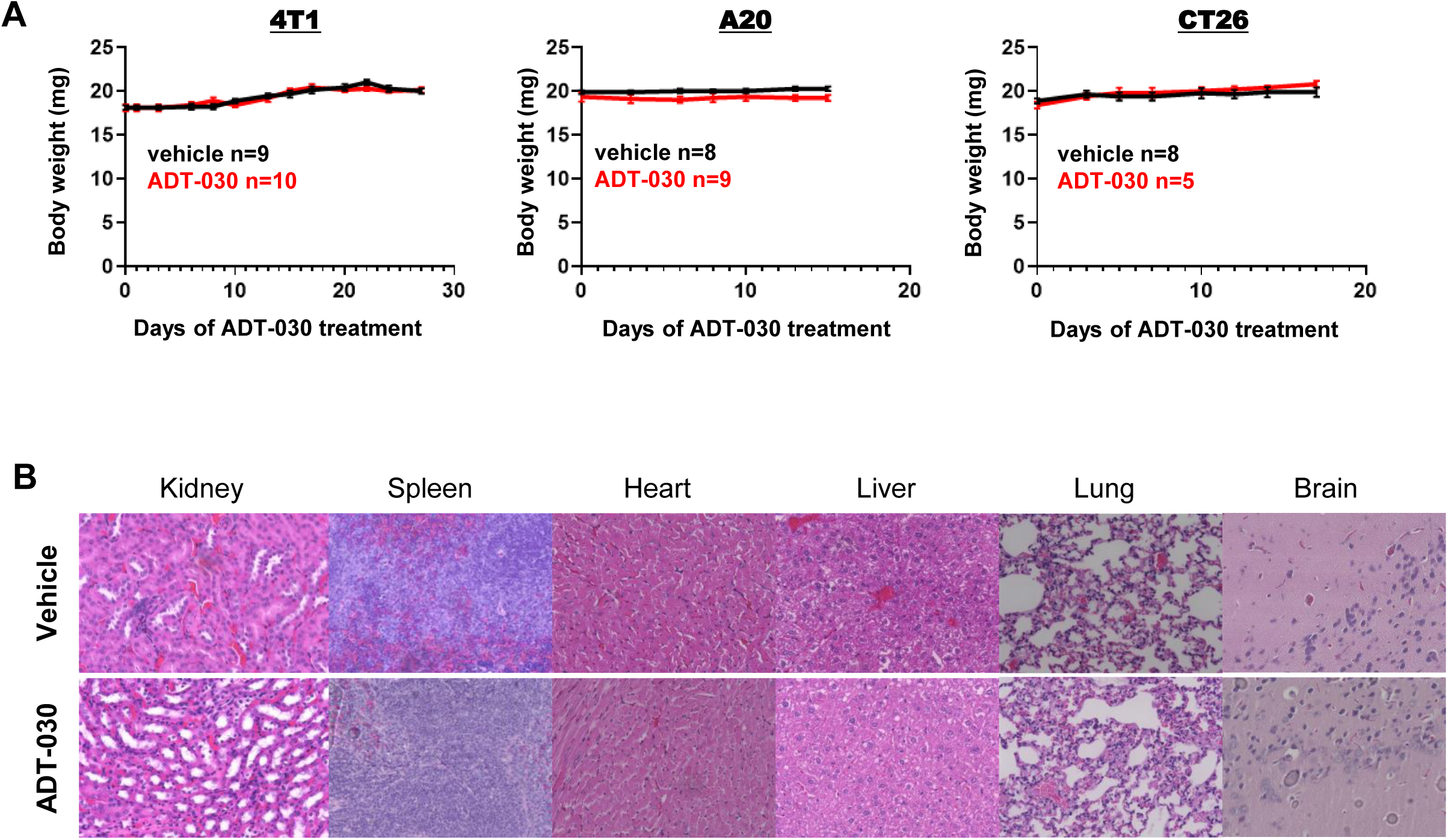
ADT-030 administration does not cause overt toxicity in mice. (A) Tumor-bearing mice were treated as described in Fig. 3A. Body weight changes over time are shown for representative tumor models and presented as mean ± SEM, with the number of mice in each group indicated. (B) Representative H/E-stained sections of the indicated organs. Naïve mice were treated with vehicle or ADT-030 for 10 days prior to tissue collection. Formalin-fixed, paraffin-embedded (FFPE) samples were processed for histological analysis..

**Supplemental Fig. 4.**
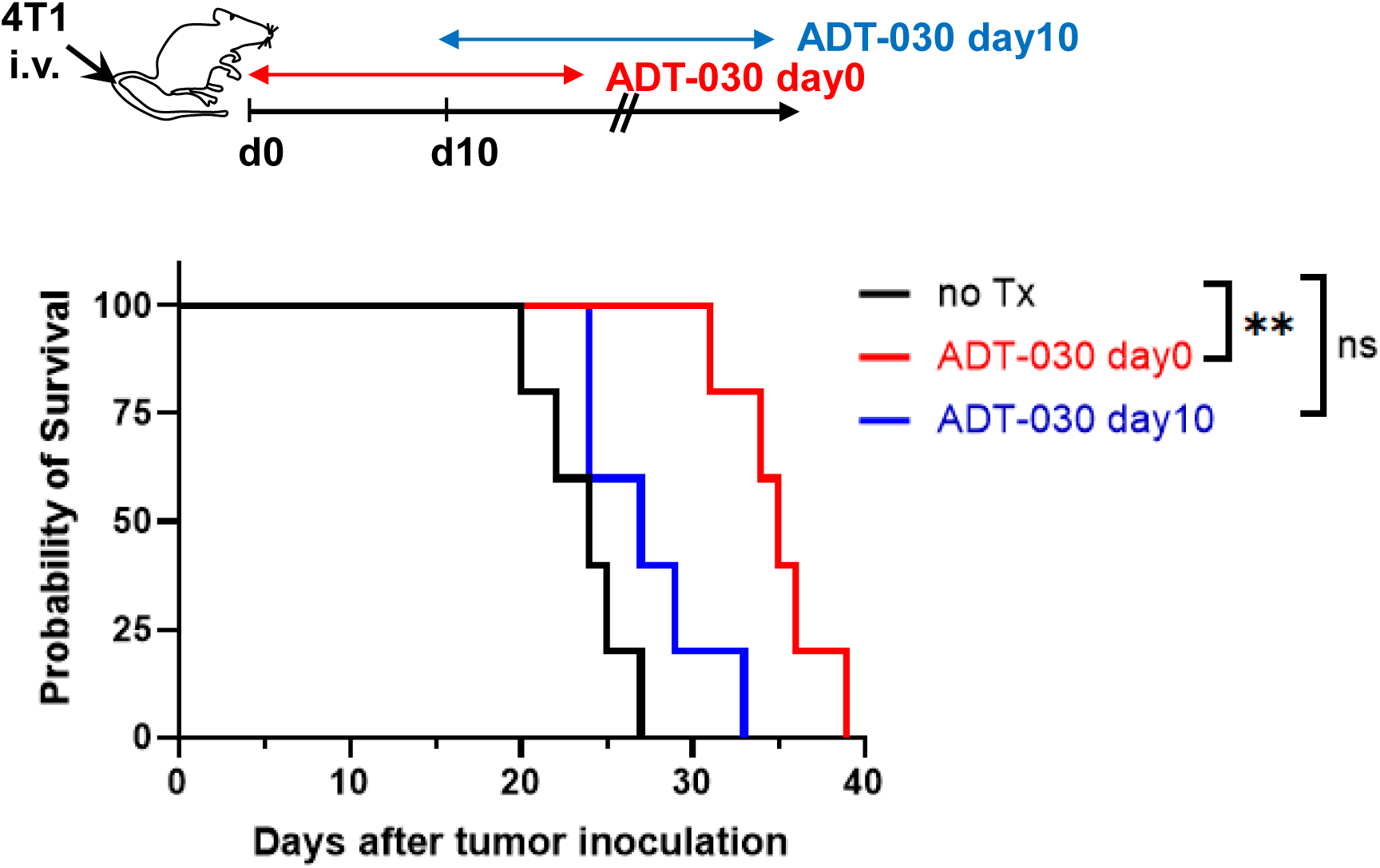
Early administration of ADT-030 provides better control of metastatic tumor growth. The schema depicts the experimental design comparing survival outcomes when ADT-030 treatment was initiated either on day 0 or day 10 after tumor inoculation. Untreated tumor-bearing mice were included as controls. Kaplan–Meier survival curves are shown, with five mice per group. ns, not significant; **, *P* < 0.01.

**Supplemental Fig. 5.**
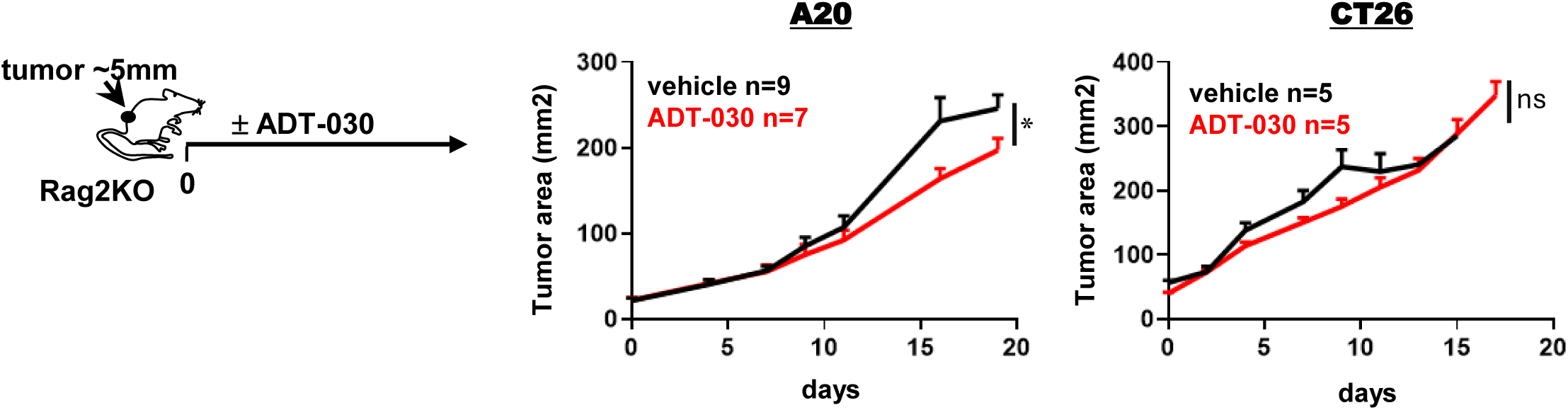
ADT-030 has diminished antitumor activity in Rag2-deficient mice. The schema depicts the experimental design. Tumor-bearing Rag2KO mice were treated as described in Fig. 5A. Data from the indicated tumor models are presented as mean ± SEM of tumor area over time, with the number of mice in each group indicated. ns, not significant; *, P<0.05.

**Supplemental Fig. 6.**
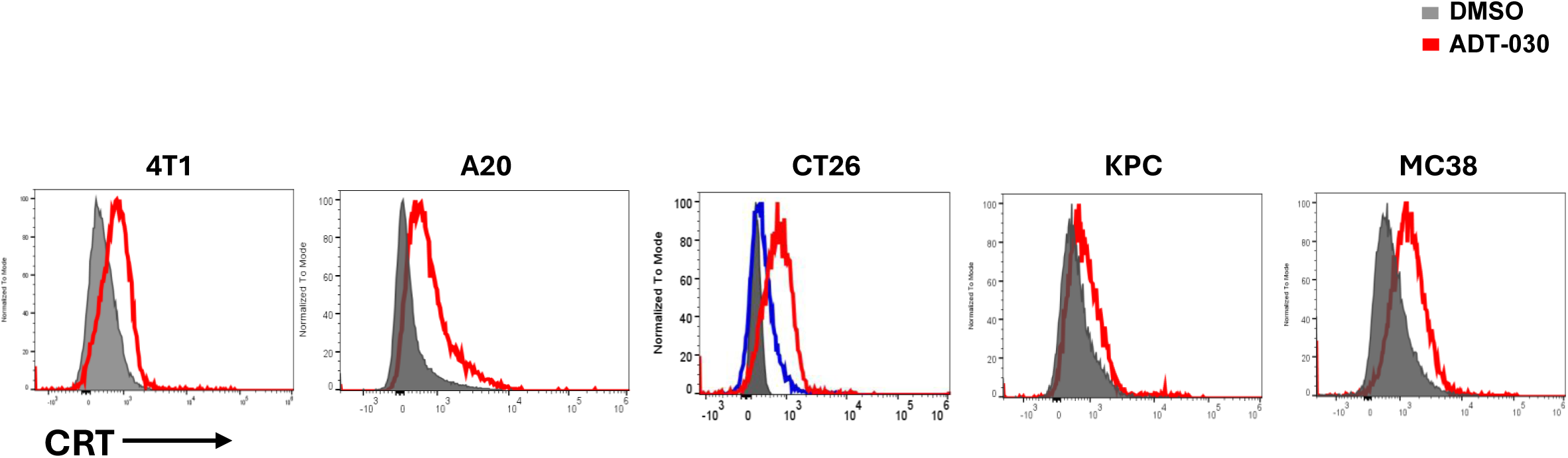
ADT-030 treatment leads to increased surface expression of CRT in tumor cells. Tumor cells were treated as described in Fig. 6A. Representative flow cytometry histograms of CRT surface expression are shown for the indicated cells.

**Supplemental Fig. 7.**
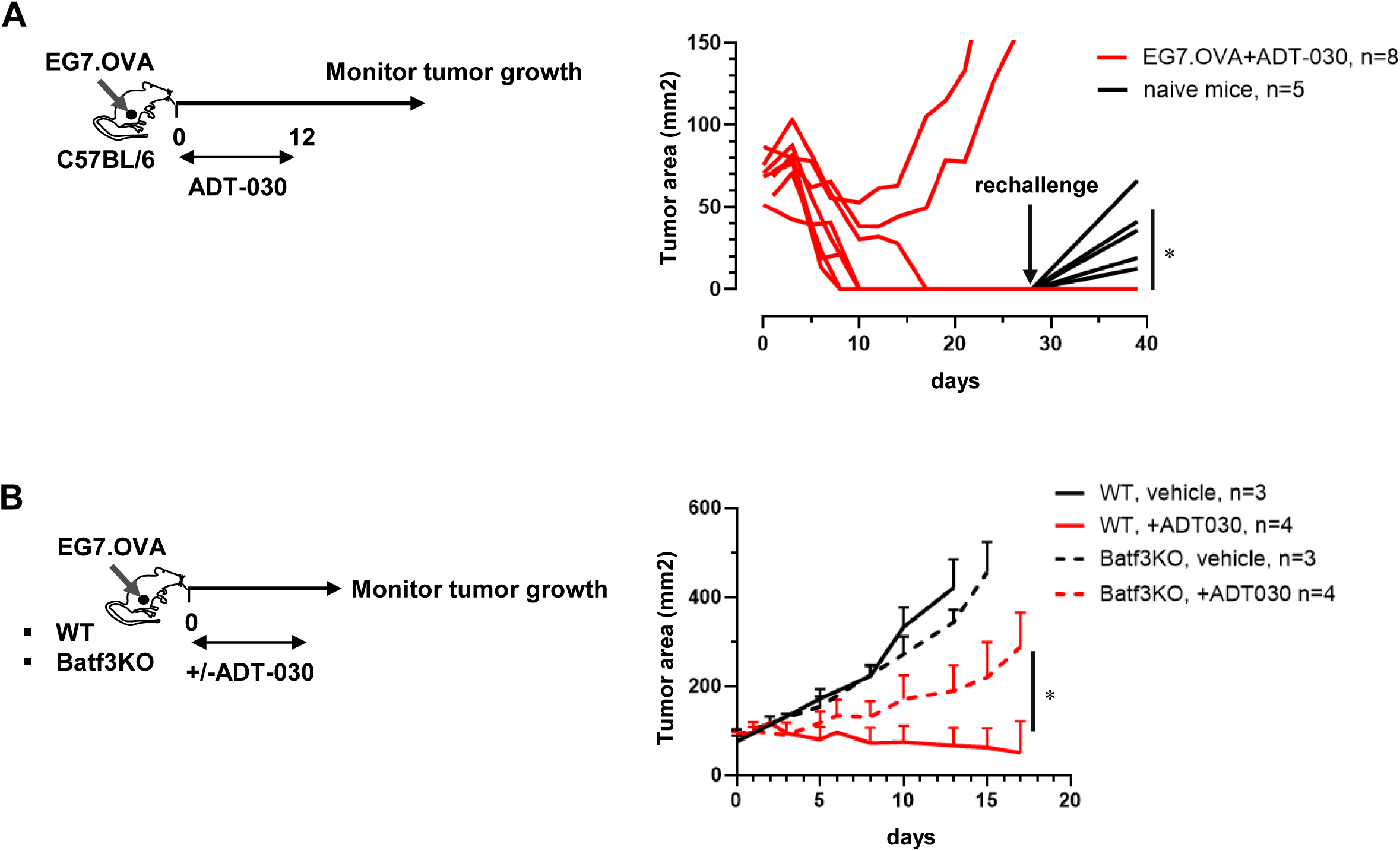
ADT-030 antitumor efficacy against EG7.OVA tumor is diminished in the absence of cDC1. (A) ADT-030 exhibits curative antitumor effect in EG7.OVA tumor model. Following the workflow depicted in the schema, WT C57BL/6 mice were implanted s.c. with EG7.OVA tumor cells. Mice with established tumors (50-100mm2) were treated with ADT-030 for 12 consecutive days. 3 mice that achieved complete tumor regression were rechallenged with EG7.OVA tumor cells. Naïve mice inoculated with EG7.OVA tumors at the same time servied as controls. (B) ADT-030 antitumor efficacy is compromised in Batf3KO mice. Following the workflow depicted in the schema, EG7.OVA cells were s.c. implanted to WT or Batf3KO mice. Mice with established tumors were treated daily with vehicle or ADT-030 daily till the endpoint. Tumor growth curves are shown as mean ± SEM, with the number of mice per group indicated. *, *P* < 0.05.

**Supplemental Fig. 8.**
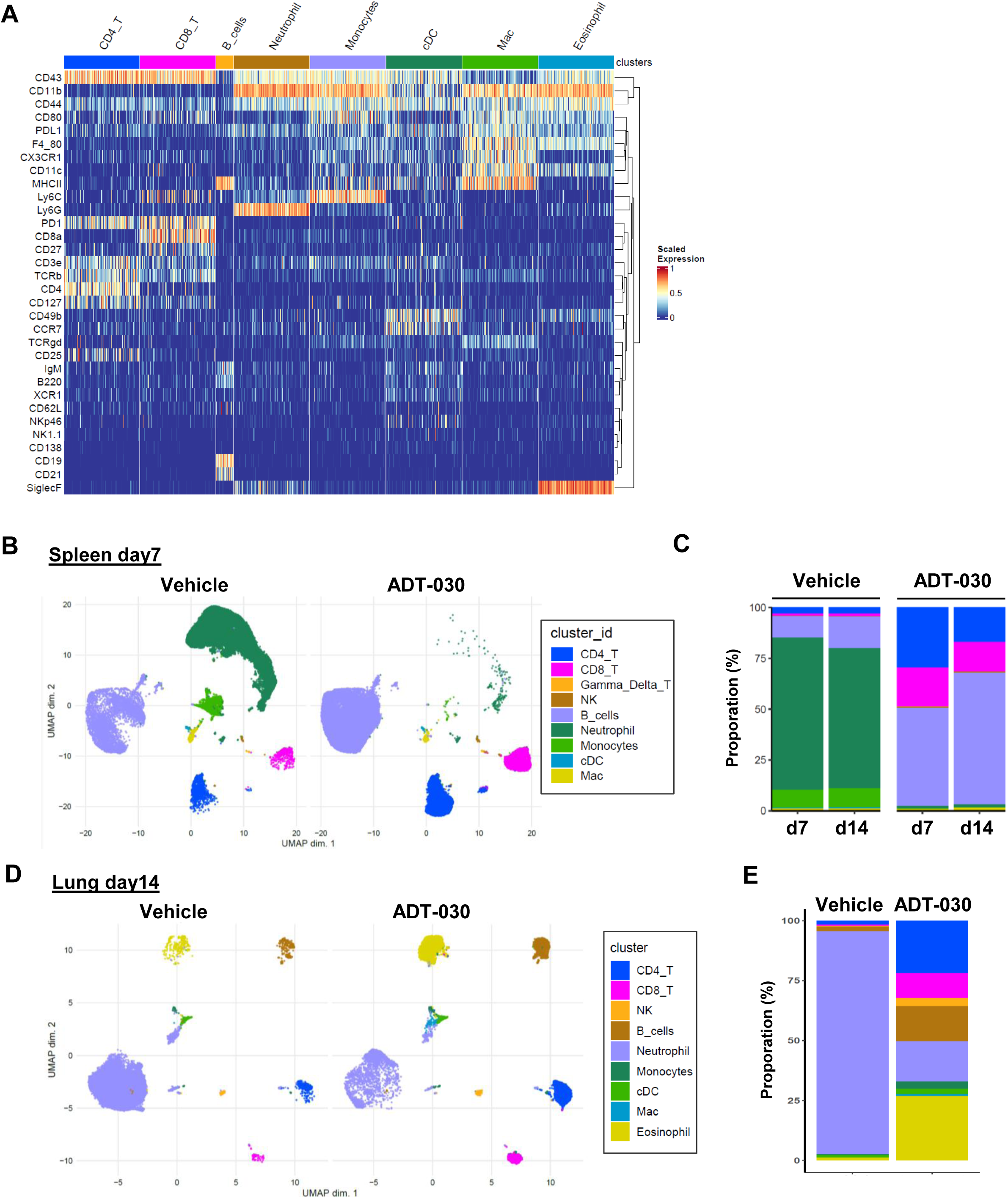
CyTOF immune profiling reveals tissue-specific immunologic changes following ADT-030 treatment. Experimental procedures for CyTOF analysis are described in Fig. 7. (A) Heatmap of scaled CyTOF marker expression at the single-cell level. (B) UMAP visualization of CD45⁺ immune cells in spleens from mice treated with vehicle or ADT-030 for 7 days. Spleens collected after 14 days were analyzed in parallel. (B) Stacked bar plots showing the relative proportions of immune cell subsets among CD45⁺ cells in spleens at different time points following the indicated treatment. (C) UMAP visualization of CD45⁺ immune cells in lungs of mice treated with vehicle or ADT-030 for 14 days. (D) Relative proportions of immune cell subsets among CD45⁺ cells in lungs.

**Supplemental Fig. 9.**
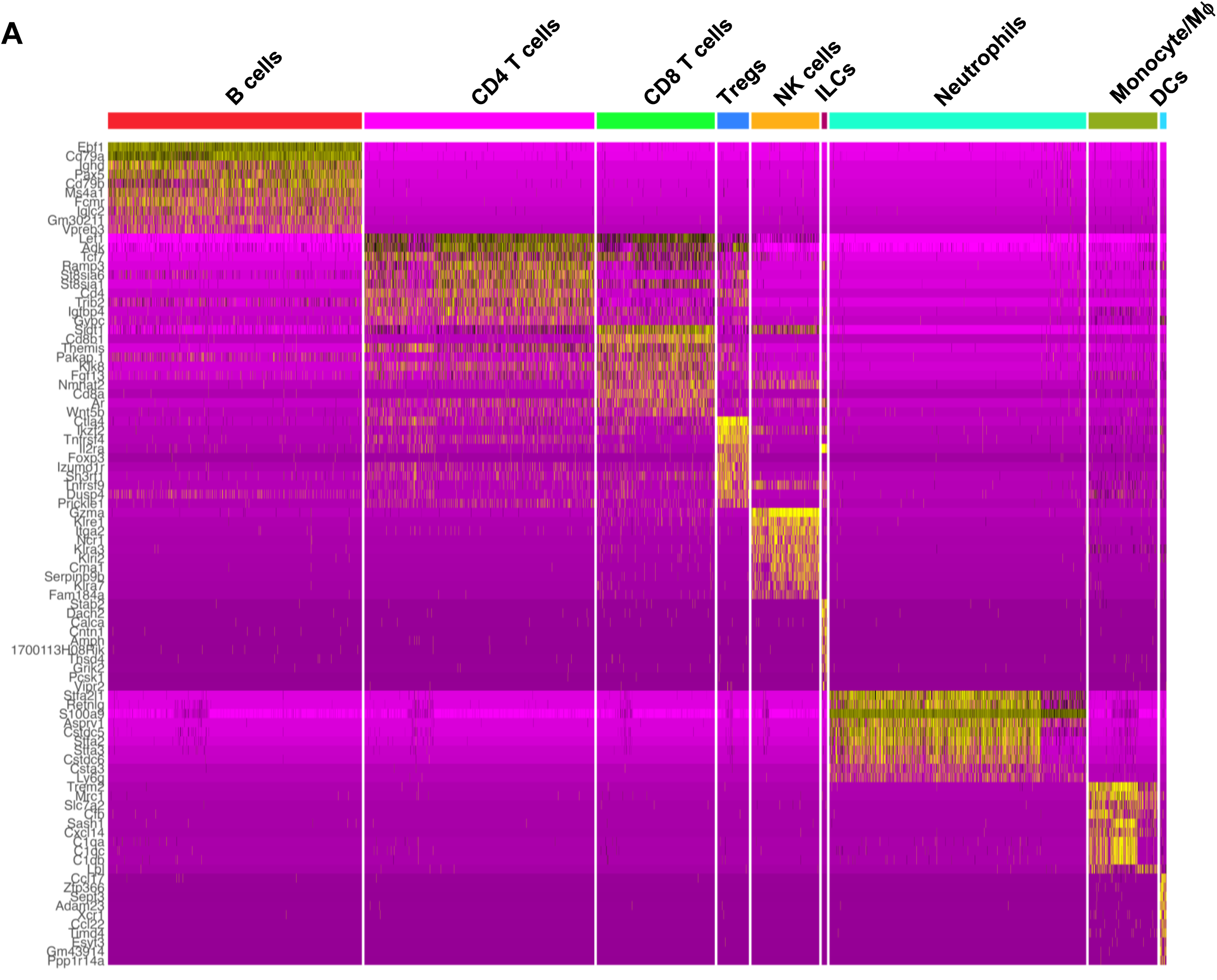
Marker gene expression defining immune cell clusters in lung scRNA-seq analysis. Heatmap showing the top 10 marker genes defining each immune cell cluster identified by scRNA-seq of CD45.1⁺ cells isolated from the lungs in the 4T1 experimental metastasis model. Columns represent individual cells and rows represent genes. Cells are grouped by cluster, with the top 10 most significant marker genes displayed for each cluster.

## References

1 Li, N. et al. Phosphodiesterase 10A: a novel target for selective inhibition of colon tumor cell growth and beta-catenin-dependent TCF transcriptional activity. Oncogene 34, 1499–1509 (2015). 10.1038/onc.2014.94

2 Zhu, B. et al. Phosphodiesterase 10A is overexpressed in lung tumor cells and inhibitors selectively suppress growth by blocking beta-catenin and MAPK signaling. Oncotarget 8, 69264–69280 (2017). 10.18632/oncotarget.20566

3 Fusco, J. P. et al. Genomic characterization of individuals presenting extreme phenotypes of high and low risk to develop tobacco-induced lung cancer. Cancer Med 7, 3474–3483 (2018). 10.1002/cam4.1500

4 Li, N. et al. Suppression of beta-catenin/TCF transcriptional activity and colon tumor cell growth by dual inhibition of PDE5 and 10. Oncotarget 6, 27403–27415 (2015). 10.18632/oncotarget.4741

5 Lee, K. et al. beta-catenin nuclear translocation in colorectal cancer cells is suppressed by PDE10A inhibition, cGMP elevation, and activation of PKG. Oncotarget 7, 5353–5365 (2016). 10.18632/oncotarget.6705

6 Lee, K. J. et al. Suppression of Colon Tumorigenesis in Mutant Apc Mice by a Novel PDE10 Inhibitor that Reduces Oncogenic beta-Catenin. Cancer Prev Res (Phila) 14, 995–1008 (2021). 10.1158/1940-6207.CAPR-21-0208

7 Borneman, R. M. et al. Phosphodiesterase 10A (PDE10A) as a novel target to suppress beta-catenin and RAS signaling in epithelial ovarian cancer. J Ovarian Res 15, 120 (2022). 10.1186/s13048-022-01050-9

8 Weijzen, S., Velders, M. P. & Kast, W. M. Modulation of the immune response and tumor growth by activated Ras. Leukemia 13, 502–513 (1999). 10.1038/sj.leu.2401367

9 Spranger, S. & Gajewski, T. F. A new paradigm for tumor immune escape: beta-catenin-driven immune exclusion. J Immunother Cancer 3, 43 (2015). 10.1186/s40425-015-0089-6

10 Yu, F. et al. Wnt/beta-catenin signaling in cancers and targeted therapies. Signal Transduct Target Ther 6, 307 (2021). 10.1038/s41392-021-00701-5

11 Bos, J. L. ras oncogenes in human cancer: a review. Cancer Res 49, 4682–4689 (1989).

12 Prior, I. A., Lewis, P. D. & Mattos, C. A comprehensive survey of Ras mutations in cancer. Cancer Res 72, 2457–2467 (2012). 10.1158/0008-5472.CAN-11-2612

13 Moon, A. Ras Signaling in Breast Cancer. Adv Exp Med Biol 1187, 81–101 (2021). 10.1007/978-981-32-9620-6_4

14 Molina-Arcas, M. & Downward, J. Exploiting the therapeutic implications of KRAS inhibition on tumor immunity. Cancer Cell 42, 338–357 (2024). 10.1016/j.ccell.2024.02.012

15 Boumelha, J., Molina-Arcas, M. & Downward, J. Facts and Hopes on RAS Inhibitors and Cancer Immunotherapy. Clin Cancer Res 29, 5012–5020 (2023). 10.1158/1078-0432.CCR-22-3655

16 Kim, D. et al. Pan-KRAS inhibitor disables oncogenic signalling and tumour growth. Nature 619, 160–166 (2023). 10.1038/s41586-023-06123-3

17 Holderfield, M. et al. Concurrent inhibition of oncogenic and wild-type RAS-GTP for cancer therapy. Nature 629, 919–926 (2024). 10.1038/s41586-024-07205-6

18 Jiang, J. et al. Translational and Therapeutic Evaluation of RAS-GTP Inhibition by RMC-6236 in RAS-Driven Cancers. Cancer Discov 14, 994–1017 (2024). 10.1158/2159-8290.CD-24-0027

19 Reddy Bandi, D. S., et al. ADT-030, a novel PDE10 inhibitor, demonstrates potent antitumor activity in pancreatic ductal adenocarcinoma. bioRxiv, 2026.2002.2011.705411 (2026). 10.64898/2026.02.11.705411

20 Foote, J. B. et al. A Pan-RAS Inhibitor with a Unique Mechanism of Action Blocks Tumor Growth and Induces Antitumor Immunity in Gastrointestinal Cancer. Cancer Res 85, 956–972 (2025). 10.1158/0008-5472.CAN-24-0323

21 Zhong, W. et al. Comparison of the molecular and cellular phenotypes of common mouse syngeneic models with human tumors. BMC Genomics 21, 2 (2020). 10.1186/s12864-019-6344-3

22 Lakics, V., Karran, E. H. & Boess, F. G. Quantitative comparison of phosphodiesterase mRNA distribution in human brain and peripheral tissues. Neuropharmacology 59, 367–374 (2010). 10.1016/j.neuropharm.2010.05.004

23 Pulaski, B. A. & Ostrand-Rosenberg, S. Mouse 4T1 breast tumor model. Curr Protoc Immunol **Chapter** 20, Unit 20 22 (2001). 10.1002/0471142735.im2002s39

24 Fucikova, J. et al. Detection of immunogenic cell death and its relevance for cancer therapy. Cell Death Dis 11, 1013 (2020). 10.1038/s41419-020-03221-2

25 Arimoto, K. I., Miyauchi, S., Liu, M. & Zhang, D. E. Emerging role of immunogenic cell death in cancer immunotherapy. Front Immunol 15, 1390263 (2024). 10.3389/fimmu.2024.1390263

26 Hildner, K. et al. Batf3 deficiency reveals a critical role for CD8alpha+ dendritic cells in cytotoxic T cell immunity. Science 322, 1097–1100 (2008). 10.1126/science.1164206

27 Ward, A. B. et al. Enhancing anticancer activity of checkpoint immunotherapy by targeting RAS. MedComm (2020) 1, 121–128 (2020). 10.1002/mco2.10

28 Youn, J. I., Nagaraj, S., Collazo, M. & Gabrilovich, D. I. Subsets of myeloid-derived suppressor cells in tumor-bearing mice. J Immunol 181, 5791–5802 (2008). 10.4049/jimmunol.181.8.5791

29 Dilly, J. et al. Mechanisms of Resistance to Oncogenic KRAS Inhibition in Pancreatic Cancer. Cancer Discov 14, 2135–2161 (2024). 10.1158/2159-8290.CD-24-0177

30 Ebright, R. Y., Dilly, J., Shaw, A. T. & Aguirre, A. J. Response and Resistance to RAS Inhibition in Cancer. Cancer Discov 15, 1325–1349 (2025). 10.1158/2159-8290.CD-25-0349

31 Ouzounova, M. et al. Monocytic and granulocytic myeloid derived suppressor cells differentially regulate spatiotemporal tumour plasticity during metastatic cascade. Nat Commun 8, 14979 (2017). 10.1038/ncomms14979

32 Gungabeesoon, J. et al. A neutrophil response linked to tumor control in immunotherapy. Cell 186, 1448–1464 e1420 (2023). 10.1016/j.cell.2023.02.032

33 Benguigui, M. et al. Interferon-stimulated neutrophils as a predictor of immunotherapy response. Cancer Cell 42, 253–265 e212 (2024). 10.1016/j.ccell.2023.12.005

34 Alicea-Torres, K. et al. Immune suppressive activity of myeloid-derived suppressor cells in cancer requires inactivation of the type I interferon pathway. Nat Commun 12, 1717 (2021). 10.1038/s41467-021-22033-2

35 Tao, Y. et al. Endogenous cGMP-dependent protein kinase reverses EGF-induced MAPK/ERK signal transduction through phosphorylation of VASP at Ser239. Oncol Lett 4, 1104–1108 (2012). 10.3892/ol.2012.851

36 Cho, K. J. et al. AMPK and Endothelial Nitric Oxide Synthase Signaling Regulates K-Ras Plasma Membrane Interactions via Cyclic GMP-Dependent Protein Kinase 2. Mol Cell Biol 36, 3086–3099 (2016). 10.1128/MCB.00365-16

37 Quadri, M. et al. Activation of cGMP-Dependent Protein Kinase Restricts Melanoma Growth and Invasion by Interfering with the EGF/EGFR Pathway. J Invest Dermatol 142, 201–211 (2022). 10.1016/j.jid.2021.06.011

38 Lan, T. et al. Increased endogenous PKG I activity attenuates EGF-induced proliferation and migration of epithelial ovarian cancer via the MAPK/ERK pathway. Cell Death Dis 14, 39 (2023). 10.1038/s41419-023-05580-y

39 Serafini, P. et al. Phosphodiesterase-5 inhibition augments endogenous antitumor immunity by reducing myeloid-derived suppressor cell function. J Exp Med 203, 2691–2702 (2006). 10.1084/jem.20061104

